# A context-dependent switch from sensing to feeling in the primate amygdala

**DOI:** 10.1101/2022.10.15.512319

**Authors:** Anne B. Martin, Michael A. Cardenas, Rose K. Andersen, Archer I. Bowman, Elizabeth A. Hillier, Sliman Bensmaia, Andrew J. Fuglevand, Katalin M. Gothard

## Abstract

The skin transmits affective signals that integrate into our social vocabulary. As the socio-affective aspects of touch are likely processed in the amygdala, we compared neural responses to social grooming and gentle airflow recorded from the amygdala and the primary somatosensory cortex of non-human primates. Neurons in the somatosensory cortex responded to both types of tactile stimuli. In the amygdala, however, neurons did not respond to individual grooming sweeps even though grooming elicited autonomic states indicative of positive affect. Instead, many showed changes in baseline firing rates that persisted throughout the grooming bout. Such baseline fluctuations were attributed to social context because the presence of the groomer alone could account for the observed changes in baseline activity. It appears, therefore, that during grooming, the amygdala stops responding to external inputs on a short time scale but remains responsive to social context (or the associated affective states) on longer time scales.

## INTRODUCTION

Gentle touch from a bonded social partner contributes to positive affect and social well-being (McGlone et al., 2014). In infancy, touch-mediated parental care stimulates brain development (Bales et al., 2018; Callaghan and Tottenham, 2016; Cascio et al., 2019), shapes the future stress-resilience of the individual (Walker et al., 2020; Dagnino-Subiabre, 2022), and lays the foundations of healthy autonomic and emotional regulation (Cadji et al., 1998; Sanchez, 2006; Gee et al., 2014). As the Romanian orphanages of the 1980s sadly demonstrated, touch deprivation in children causes irreversible emotional and social-cognitive deficits (Rutter, 1998; O’Connor and Rutter, 2000; Mackes et al., 2020). Even after infancy and childhood, communication through touch remains fully integrated into our social vocabulary, allowing us to understand a rich variety of emotional signals through the skin (Hertenstein et al., 2006; McGlone et al. 2014). In humans and non-human primates, socially appropriate affective touch between adults builds long-lasting, trusting bonds (Von Mohr et al., 2017; Dunbar 2010, 2022).

Grooming in macaques is the equivalent of social and affective touch in humans. Beyond its hygienic role, grooming maintains the social homeostasis of hierarchical societies (Schino et al., 1988; Lehmann et al., 2007; Schino and Aureli, 2008) and benefits both the groomer and the recipient. The groomer builds alliances and gains coalition support, tolerance at feeding sites, infant handling, and even a potential rise in the hierarchy (Dunbar, 2010; Tiddi et al., 2012; McFarland and Majolo, 2011). The recipient attains a physiological state marked by muscle relaxation, reduced anxiety and vigilance, enhanced vagal tone, and the release of oxytocin and endorphins that counter the effects of circulating glucocorticoids induced by previous stressors (Boccia et al.,1988; Aureli et al., 1999; Grandi and Ishida 2015; Jablonski, 2021). Similar physiological benefits have been documented in humans who receive affective touch from bonded partners (Walker et al., 2020; Dagnino-Subiabre, 2022; Triscoli et al., 2017; Moberg and Petersson, 2022; Nummenmaa et al., 2016), and in macaques who are groomed by trusted human caregivers (Taira and Rolls, 1992; Grandi and Ishida, 2015). These observations led to the prediction that neurons in the amygdala would respond differentially to grooming and to innocuous tactile stimuli delivered to the same areas of the skin.

Electrophysiological studies in macaques have shown that the amygdala responds robustly to somatosensory stimulation (Livneh et al., 2012, Mosher et al., 2016, Morrow et al., 2019) although often integrated with other sensory modalities, task variables, actions required, and even abstract features such as behavioral context (Gothard, 2020). This is likely a consequence of convergence of multiple circuits in the amygdala that process sensory, affective, social, and autonomic signals (Amaral, 1992; LeDoux, 2007; Bickart et al., 2014). By virtue of its connections, the amygdala is in an ideal position to extract the affective significance of tactile stimuli and to enlist autonomic effectors to generate the corresponding physiological states (Gothard and Fuglevand, 2022). Indeed, the human amygdala is activated by pleasant touch (Löken et al., 2011; Lukas et al., 2014; Suvilehto et al., 2021). The magnitude of this activation depends not only on the mechanical properties of the stimuli, the activation of C-tactile afferents (Olausson et al., 2010), but also on the body part touched and the relationship between the recipient and the deliverer of tactile stimulation (Suvilehto et al., 2021). Affective touch also enhances the functional connectivity of the amygdala with several other cortical areas, including the subdivision of medial prefrontal cortex (Gordon et al., 2013, Rolls et al., 2003) involved in affective and social processes (Ghangophaday et al., 2021).

Given the responsivity of the amygdala to tactile stimulation and its processing bias for stimuli with affective significance, we asked whether neurons in the amygdala would respond differentially to grooming and to an innocuous tactile stimulus. Specifically, we compared the tactile responses elicited by gentle airflow to responses elicited by grooming-like finger sweeps delivered by a trusted human trainer. We also recorded responses to these two forms of tactile stimuli at an early cortical processing stage in the primary somatosensory cortex (SI, Brodmann’s area 3b; Kaas, 1983). Airflow elicited robust, responses from both areas. Surprisingly, however, episodic responses to grooming in the amygdala were absent. Furthermore, under different social contexts accompanied by disparate autonomic states, long-lasting modulation of baseline activity in the amygdala was observed. This suggests that the amygdala switches into different modes of sensory processing depending on the social situation. The transmutation of sensing to feeling, therefore, may not only depend on the type of skin mechanoreceptors activated by touch (McGlone et al., 2014), but also on the instantiation of social context by the amygdala.

## RESULTS

### A naturalistic design to compare neural responses elicited by affective and neutral touch

Grooming-like tactile stimuli were delivered to each subject by a trusted human trainer who emulated the natural grooming movements in monkeys through gentle, repeated sweeps of the index finger across different regions of the face (Fig. 1A). These grooming-like stimuli were applied primarily to two face regions, the upper muzzle and the brow contralateral to the recording electrodes. The trainer delivered 10 repeated sweeps (each sweep ∼ 1 s in duration) to the same location on the face before moving to a different face location, approximating the pattern of natural grooming.

**Figure 1.**
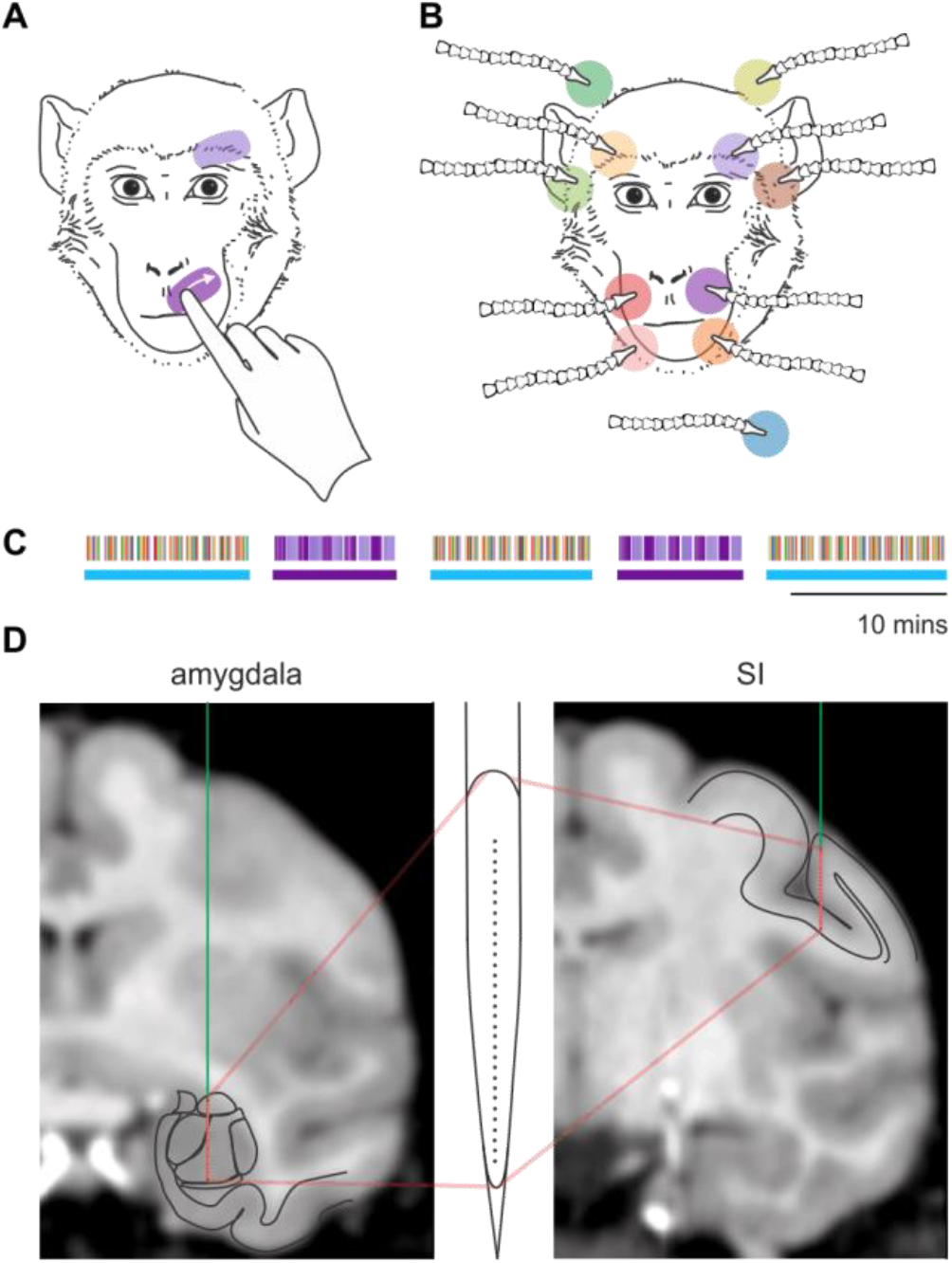
Experimental design. **A**. Two areas of the face that received grooming sweeps. **B**. Ten areas of the face that received airflow stimuli. One nozzle (pale blue disk) served as sham. **C**. Example set of alternating blocks of airflow and grooming stimulation, with block type indicated by horizontal blue and purple bars, respectively. Thin vertical lines of different color indicate the sequence of stimulation sites, color-coded according to the locations shown in panels A and B. **D**. Example recording sites in the amygdala (left) and SI (right). The V-probe line drawing shows that the 32 contacts spanned the ∼ 6 mm of the vertical axis of the monkey amygdala.

The grooming sweeps were contrasted to gentle, non-startling airflow (1 s duration) delivered through a set of adjustable nozzles directed to 10 distinct regions of the face plus a sham airflow directed away from the face (Fig. 1B). Our previous work showed that neurons in the amygdala that respond to tactile stimulation of the face have broad receptive fields, sometimes bilateral or ipsilateral to the recorded amygdala (Mosher et al., 2016; Morrow et al., 2019). This arrangement of nozzles ensured reasonable coverage of the face and seemed likely to elicit tactile responses in a subset of recorded neurons. During airflow stimulation, the monkey sat alone in the recording booth. For practical purposes, only two of the ten areas that received airflow were groomed.

Blocks of grooming alternated with blocks of airflow (Fig. 1C). For grooming blocks, 10 grooming sweeps were repeated five times for each face area, with a duration of ∼ 10 minutes. For airflow blocks, stimuli were delivered through the 11 nozzles in a random sequence and were repeated 10 times within a block. The total duration of an airflow block was about 12 minutes. Although the pattern of airflow blocks interspersed with grooming was generally upheld, the sequence and order of blocks was altered in about 1/3 of the sessions to accommodate various controls.

We used two 32-channel V-probes to record neural responses to the two types of tactile stimuli in area 3b of the primary somatosensory cortex (SI) and from the amygdala simultaneously. Of the 315 neurons in recorded in SI, 269 responded to at least one tactile stimulus. Of the 615 neurons recorded in the amygdala, that had an average firing rate >1Hz and were stable for a sufficient number of trials to assess stimulus responsiveness (see Methods), 333 responded to a tactile stimulus. As expected, neurons in SI responded differentially to the two types of stimuli and showed spatial selectivity. In SI, responses to airflow exhibited a strong transient component at the onset of the stimulus followed by a weaker, sustained component that lasted for the duration of the stimulus (example neuron, Fig. 2A). On the other hand, responses to grooming exhibited a gradual increase and decrease in firing rate (Fig. 2B), tracking the time course of pressure applied by the finger (see Fig. 3A). While none of the SI neurons responded to the sham stimulus (airflow nozzle directed away from the monkey), 94 of the 333 tactile-responsive amygdala neurons did respond to the sham. This was not surprising because a large proportion of neurons in the amygdala are multisensory and might have responded to the auditory component of the airflow (Morrow et al., 2019).

**Figure 2.**
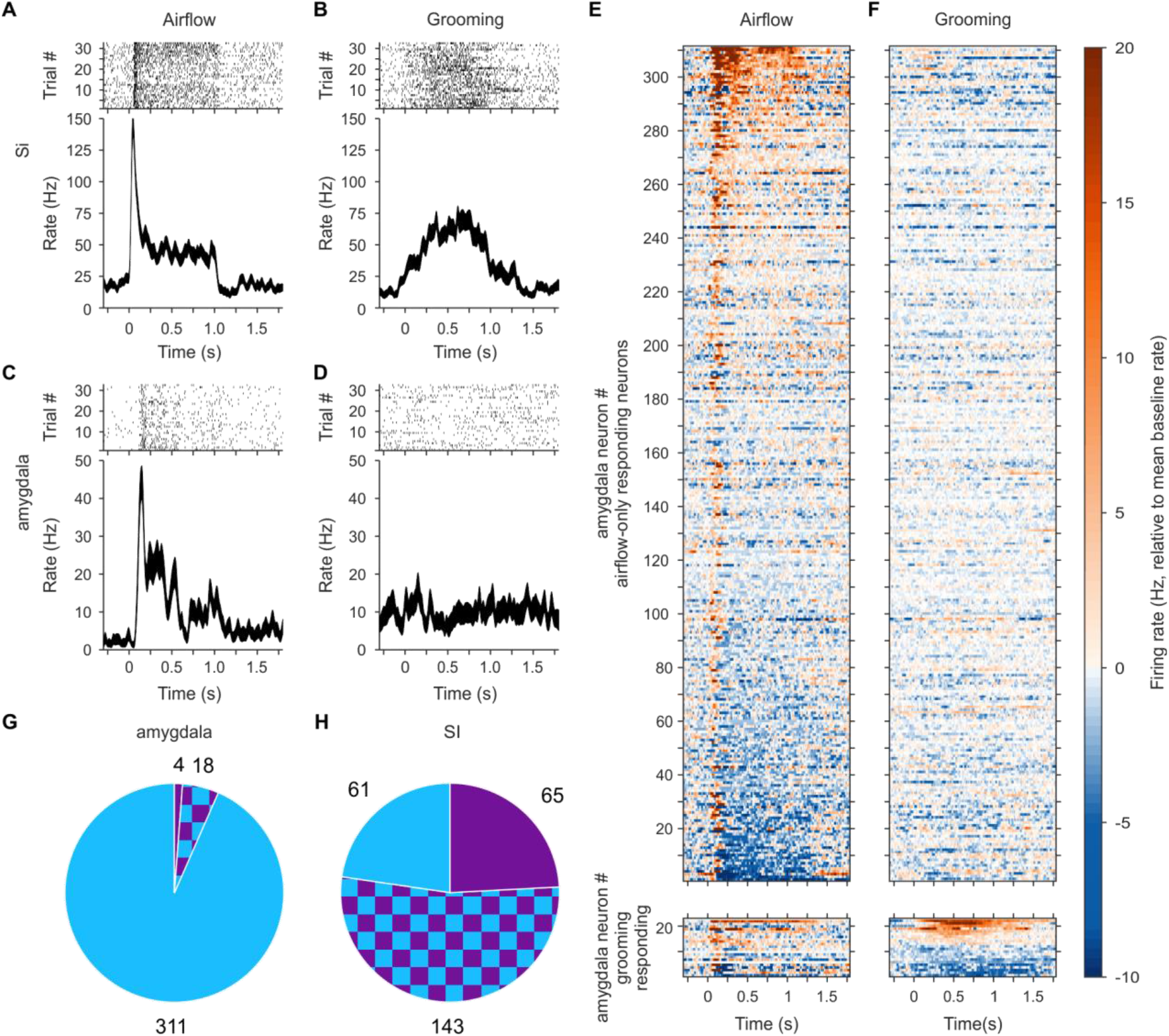
Responses to airflow and grooming stimuli in SI and the amygdala. **A, B**. Example SI neuron that responded to both airflow (**A**) and grooming (**B**) stimuli. Raster plots (top) and spike density function ± sem (bottom) for stimuli targeting the left upper muzzle, aligned to airflow or grooming sweep onset. **C, D**. Example amygdala neuron that responded only to airflow stimuli. **E, F**. Population raster of tactile-responsive neurons in the amygdala depicting the mean activity relative to baseline, aligned to airflow (**E**) or grooming stimuli (**F**). Neurons responding only to airflow (n = 311, top) are sorted by the strength of their airflow response. Neurons with grooming responses (n = 22, bottom) are sorted by the strength of their grooming response. **G, H**. The relative proportion of tactile-responsive neurons recorded from the amygdala (**G**) and SI (**H**), with airflow-only responses (blue), grooming-only (purple), and both (checkered).

**Figure 3.**
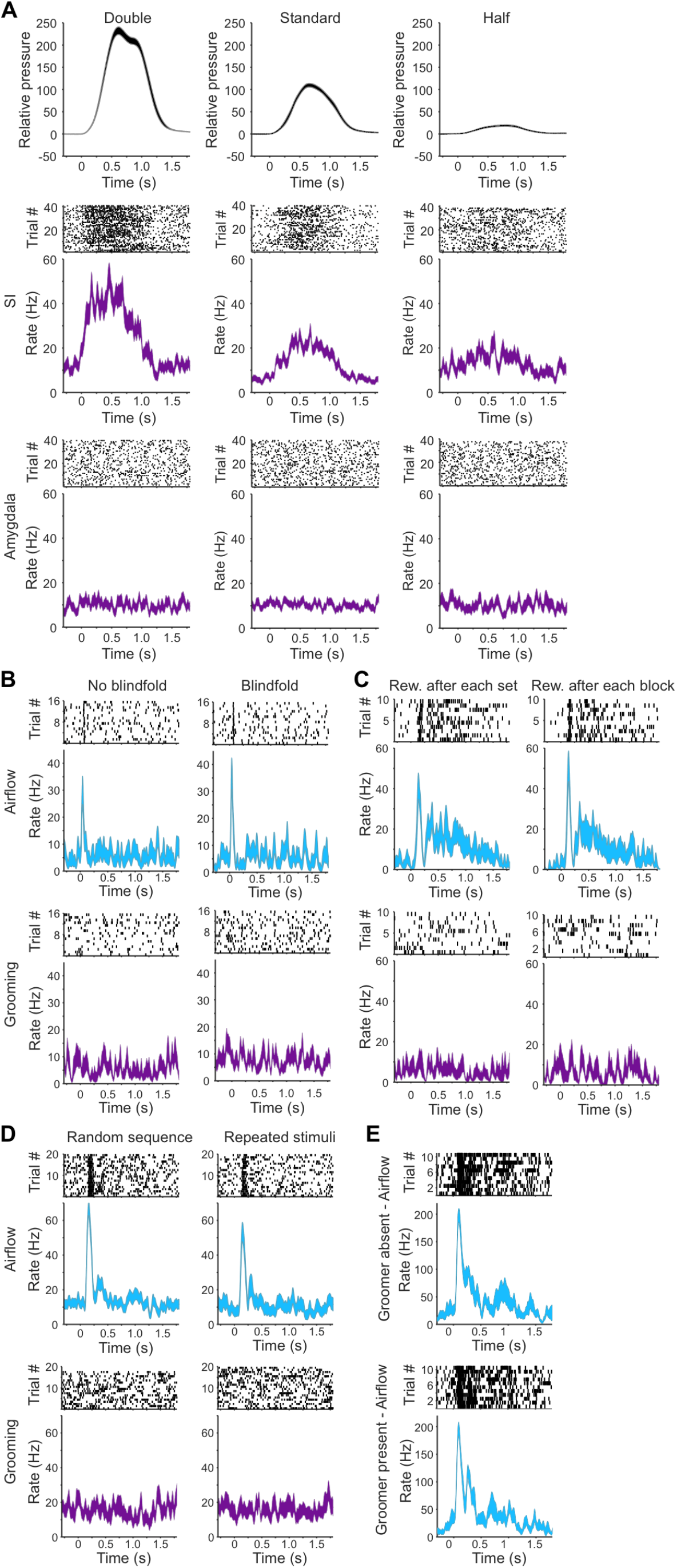
Responses of individual neurons from control experiments. **A**. Responses of example neurons to altered grooming for higher (twice standard, left panels), standard (center panels), and lower (half standard, right panels) pressures. Top panels show mean contact pressure ± sem. Middle panels show the activity of an example responsive SI neuron. Bottom panels show activity of an example non-responsive amygdala neuron. Raster plots (above) and spike density functions ± sem (below) are aligned to grooming sweeps targeting the left upper muzzle. **B**. Responses of an example amygdala neuron to airflow (top) and grooming (below) without (left) and with blindfold (right). **C**. Responses of an example amygdala neuron to altered reward contingencies. Reward after each set (left) is the standard protocol for airflow. Reward after each block (right) is the standard protocol for grooming. Responses to airflow (top) and grooming (below). **D**. Responses of an example amygdala neuron to altering the sequence of airflow (top) and grooming stimuli (below). Random sequence (left) is the standard protocol for airflow. Repeated stimuli (right) is the standard protocol for grooming. **E**. Responses of an example neuron to airflow when the groomer was absent (top) and present (bottom).

### Suppression of neural responses to individual grooming sweeps in the amygdala

Compared to neurons in SI, neurons in the amygdala exhibited a different pattern of responses to tactile stimuli. As shown in Fig. 2C, the example amygdala neuron exhibited a strong transient at airflow onset, which was followed by a weaker, uneven response. Surprisingly, however, grooming stimuli applied to the same region of the face failed to evoke a response in this neuron (Fig. 2D). This pattern of responsivity to airflow (Fig. 2E) but not to grooming (Fig 2F) was observed in most amygdala neurons. Indeed, most of these neurons responded exclusively to airflow (311/333, 93%) (Fig. 2G), and only a small fraction (4/333, 1.2%) were exclusively activated by grooming. This outcome was at odds with our expectation that in the tactile domain, as in the visual domain, neurons in the amygdala would show strong responses to the stimuli with high social and affective significance. In comparison, the majority (53%) of the 269 tactile-responsive neurons in SI responded to both grooming and airflow (Fig. 2H). Because of well delineated and small receptive fields in the face region of area 3B in SI (Nelson et al. 1980), slight differences in the locations of airflow and grooming stimuli may partially account for the lower percentage than expected of neurons responding to both types of tactile stimuli. About 23% of neurons responded exclusively to airflow and another 24% to grooming sweeps applied to the face. Overall, 77% of tactile-responsive neurons in SI responded to grooming, while only 7% did so in the amygdala.

### Control Experiments

To establish that the observed difference between SI and the amygdala in response to grooming was not explained solely by differences in the mechanical properties of stimuli or other features of experimental design, we carried out five control experiments (Fig. 3).

First, we tested whether suppressed responses to grooming were due to inadequate stimulus pressure. To do this, we measured neural responses in the amygdala and SI to grooming stimuli that were approximately doubled and halved in magnitude compared to the standard grooming pressures (Fig. 3A). While SI neurons were clearly modulated by grooming magnitude, only 1/18 amygdala neurons recorded in these experiments responded to grooming at any pressure.

Second, the visual component of receiving grooming (i.e., the hand and finger of the experimenter looming toward the groom sites on the monkey’s face) might have led to differential responses in the amygdala to the two forms of tactile stimuli. To control for this factor, we compared responses to grooming with and without blindfolding the animal. As shown in the example neuron in Fig. 3B, blindfolding did not alter its responses to airflow or grooming stimuli. Collectively, blindfolding had no effect on number of neurons in the amygdala responsive to airflow or grooming stimuli (chi-square *p* = 0.99, n = 40).

Third, we explored the possibility that reward contingencies used in the original experiments might have contributed to the observed results. In the initial design, reward was delivered after each sequence of 11 airflow stimuli. During grooming blocks, however, the animal received reward only after the block was completed. This led to substantially different intervals of time between rewards that could have altered the animal’s reward expectations and potentially altered responsiveness in the amygdala. Therefore, we inverted the reward scheme by providing reward after each set of grooming stimuli and provided reward only at the end of airflow blocks. This change had no effect on the observed neural responses in the amygdala (Fig. 3C). Overall, there was no change in the proportion of amygdala neurons responsive to grooming or airflow with alteration in reward contingencies (chi-square *p* = 0.7, n = 40).

Fourth, we varied the predictability of stimuli, either delivering them randomly or in a particular sequence. In the standard protocol, airflow stimuli were delivered to 10 different areas of the face in an unpredictable random sequence while grooming sweeps were delivered to two locations, ten consecutive times at each location. This grooming protocol, therefore, was predictable, perhaps increasing the likelihood of habituation and potentially reducing error prediction. For these control experiments, the pattern of delivery was reversed: airflow stimuli were delivered to the same two locations in groups of 10 sequential stimuli (as was done for standard grooming sweeps), whereas grooming sweeps were delivered to 10 different locations in a random sequence (as done for standard airflow stimuli). The time between repeated airflow stimuli was precisely 3 s, enhancing their predictability, whereas the timing between grooming stimuli was made variable, augmenting their unpredictability. These alterations in the sequencing and the predictability of stimuli did not change the responsiveness of amygdala neurons to airflow or to grooming (Fig. 3D, airflow chi-square *p* = 0.2, n = 36; no neurons responded to grooming with either predictable or unpredictable stimuli).

And fifth, we asked whether the presence of the groomer (rather than grooming per se) led to suppressed neural responses. For these experiments, the groomer was present in the recording booth during blocks of trials involving airflow stimuli. We did not find a difference in the responsiveness of amygdala neurons to airflow, whether the groomer was present or absent during airflow blocks (Fig. 3E, chi-square test *p* = 0.99, n = 54).

Across the 188 amygdala neurons tested in all five controls, we did not observe systematic changes in the proportion or in the strength of responses to tactile stimuli (Fig. S1). The results of these control experiments suggest that the suppression of grooming responses in the amygdala was not a consequence of the above five specific features of the experimental design.

### Sustained changes in the baseline activity may transmit information about social context

While carrying out these experiments, we observed that the *baseline* firing rates appeared different in the airflow and grooming blocks in some neurons. For example, the baseline firing rate of the neuron shown in Fig. 2D was higher before the grooming stimulus was applied (∼ 10 Hz) than before the airflow stimulus was applied (∼ 2Hz, Fig. 2C). We therefore examined changes in baseline activity (binned into 1-s epochs) of each neuron for each recording session by concatenating segments of baseline activity that fell between stimuli (excluding 300 ms before and after stimulus delivery). For this analysis, we used only a subset of 237 neurons that showed stable firing rates across four consecutive blocks alternating between airflow and grooming.

To determine whether individual neurons exhibited systematic shifts in baseline firing rates, we calculated the effect size (Cohen’s *d*_*s*_: difference in mean baseline firing rates divided by the pooled standard deviation, Lakens 2013) across airflow and grooming blocks. We used this approach, rather than standard statistical inference testing (e.g., t-tests), because of the high probability of false positives when using large sample sizes (Granger 1998; Lin et al. 2013; Krzywinski & Altman 2014; Nuzzo 2014), as we had many 1-s samples distributed over sessions often lasting an hour or longer. Furthermore, to help ensure that detected changes were not due, for example, to progressive changes in baseline over the course of an experiment, we also calculated the effect sizes across blocks of the same condition (i.e., airflow block 1 vs. airflow block 2 and grooming block 1 vs. grooming block 2). For a neuron to be considered as showing context-related changes in baseline firing rate, the grooming-to-airflow effect size needed to exceed the standard minimum of 0.2 (Cohen, 1992) and be 1.5 times greater than the largest effect size of the within condition measures (see Fig. S2).

Under these criteria, 60 neurons were identified as showing clear context-related baseline activity. Of these 60, 40 had baseline firing rates greater during grooming than during airflow (Fig. 4A), whereas 20 had baseline firing rates greater during airflow than grooming (Fig. 4C). Collectively, these 60 cells had a median grooming to airflow *d*_*s*_ = 0.49 (min = 0.2, max = 1.9). Fig. 4D shows an example neuron that increased its baseline firing rate during both grooming blocks and remained at a lower level during both airflow blocks. Conversely, Fig. 4H shows and example neuron that decreased its baseline firing rate during both grooming blocks and remained at higher levels during both airflow blocks. One hundred seventy-seven neurons (Fig. 4B) did not fulfill the criteria for context-related activity in their baseline firing rates, although many of these neurons showed similar patterns activity to the neurons shown in Fig 4D. For example, the neuron shown in Figure 4E had higher baseline rates during grooming compared to airflow but failed our strict criteria, primarily because of systematic variation in baseline firing rates across the two grooming blocks. Figure 4F depicts a neuron with little systematic variation in baseline across different blocks. The neuron shown in Figure 4G had higher baseline rates during airflow than grooming but also failed the criteria because of modulation in baseline rates across the first two airflow blocks. Collectively, ∼25% of the recorded neurons in the amygdala exhibited baseline activities that seemed to be influenced by context.

**Figure 4.**
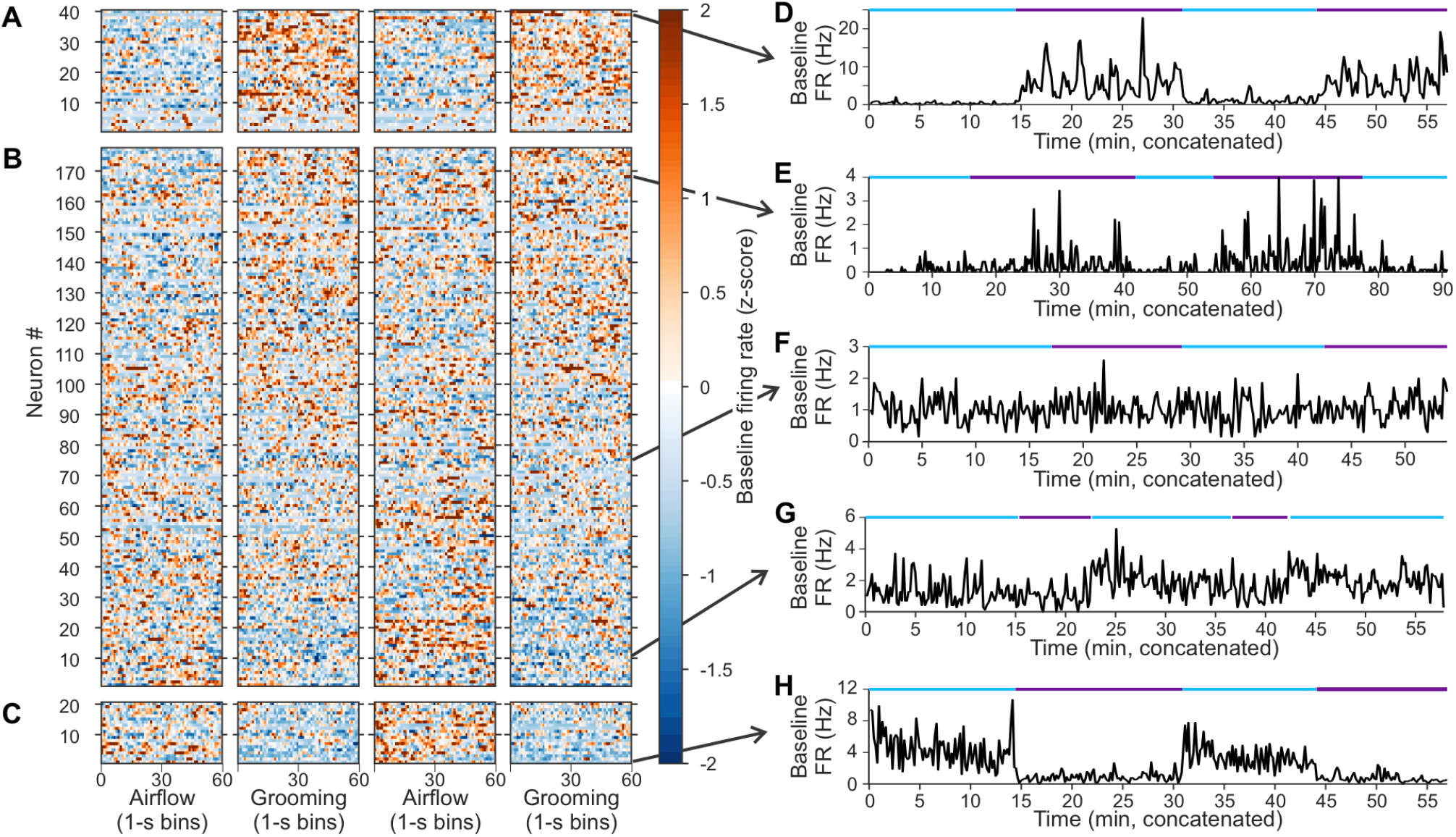
Context-related modulation of baseline firing rates. **A**. Baseline firing rates of 40 neurons identified as exhibiting context-related baseline activities for which baseline firing rate was greater during grooming than airflow. Firing rates (1-s bins) are represented as a Z-score. Bins were uniformly sampled throughout the block (e.g., if a block was 600 s in duration, then every 10^th^ bin was selected for plotting). Neurons are sorted from smallest to largest effect size for the grooming-airflow comparison. **B**. The set of 177 neurons that did not pass the criteria for exhibiting context-related activity in baseline firing. Neurons are sorted from those having the most negative effect size (i.e., airflow > grooming) to those having the greatest positive effect size (i.e., grooming > airflow). **C**. The set of 20 neurons identified as exhibiting context-related baseline activities for which baseline firing rate was greater during airflow than grooming. **D**. Example neuron with elevated baseline firing rate during grooming relative to airflow (effect size *d*_*s*_ = 1.85). Firing rates smoothed with a 10-s Gaussian filter. Blue and purple lines indicate periods of airflow and grooming, respectively. Arrow indicates position in the raster in panels A – C. **E**. Example neuron that did not pass the criteria as showing context-related activity. Nevertheless, baseline firing rate was higher during grooming compared to airflow (*d*_*s*_ = 0.56). **F**. Example neuron with minimal modulation in baseline firing rate across airflow and grooming blocks (*d*_*s*_ = -0.03). **G**. Example neuron that failed to meet the criteria as having context-related baseline activity, although it possessed greater firing rate during airflow compared to baseline (*d*_*s*_ = -0.55). **H**. Example neuron with clearly elevated baseline firing rate during airflow compared to grooming (*d*_*s*_ = -0.99).

To explore whether such context-related activity might reflect the differences in the social component of the two testing situations, we carried out additional experiments. We recorded 54 neurons while the groomer simply sat quietly in the recording booth during presentation of airflow stimuli. We compared the baseline activities of the neurons when the groomer was present and absent during airflow stimuli. When the groomer was present, baseline activity showed small but significant changes compared to airflow blocks without the groomer. Cells that showed increased baseline firing rates during grooming increased their baseline firing rate when the groomer was present during the airflow blocks (white bar in Fig. 5A) (mean increase ± SD, 0.48 ± 0.14 Hz increase, n = 29 cells, paired t-test p < 0.001). Likewise, for cells that that showed decreased baseline firing rates during grooming, the presence of the groomer reduced baseline firing rate during airflow (Fig. 5B) (mean decrease = 0.64 ± 0.2 Hz, n = 25 cells, *p* < 0.01).

**Figure 5.**
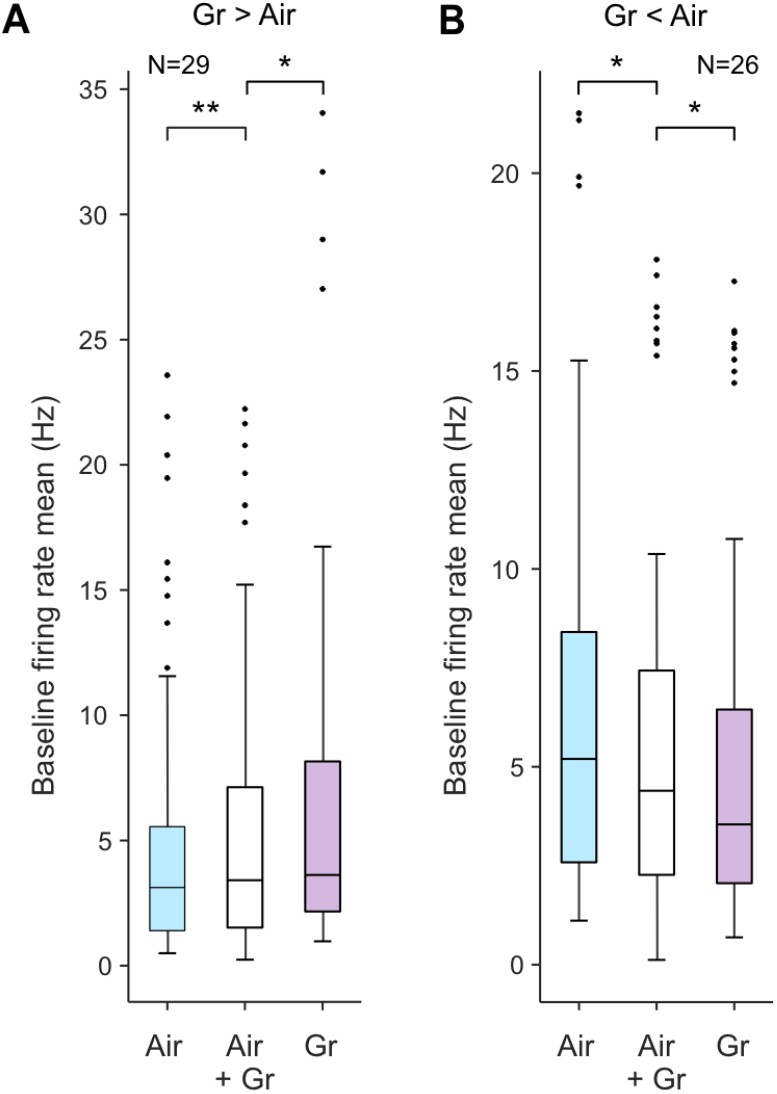
Effect of groomer presence on baseline firing rate. **A**. Average baseline firing rate of 29 neurons with enhanced rates during grooming relative to airflow across three conditions: standard airflow with the groomer absent (blue bar), airflow blocks with the groomer present (white bar), and standard grooming blocks (purple bar). * indicates p < 0.01, ** p < 0.001. **B**. Average baseline firing rate of 26 neurons with decreased baseline rates during grooming relative to airflow, across the same three conditions.

Even though the presence of the groomer was sufficient to partially induce these changes, grooming further enhanced or decreased the baseline firing rates of these neurons relative to the airflow blocks (Fig. 5A, 0.84 ± 0.24 Hz increase relative to airflow with groomer present, t-test p < 0.01; Fig, 5B, 0.44 ± 0.20 Hz, p < 0.01). This pattern of gradual shift suggests that the presence of the groomer and overt grooming stimuli might be additive in representing social context.

To explore more systematically the putative context representation through baseline firing rates, we determined whether a linear classifier (a support vector machine, SVM) could accurately decode context from the baseline firing rates of amygdala neurons. We first determined the performance of the SVM for each of the 237 neurons using 10-fold cross-validation on 1-s bins of baseline firing rates (Fig. 6A). We found that the baseline activity of 127 individual neurons were predictive of context. For each of these neurons, the classifier correctly assigned more time bins to airflow or grooming blocks than would be expected by chance (i.e., the mean performance across folds was greater than the 95% confidence interval of the null distribution computed for each neuron, gray shading in Figure 6A).

**Figure 6.**
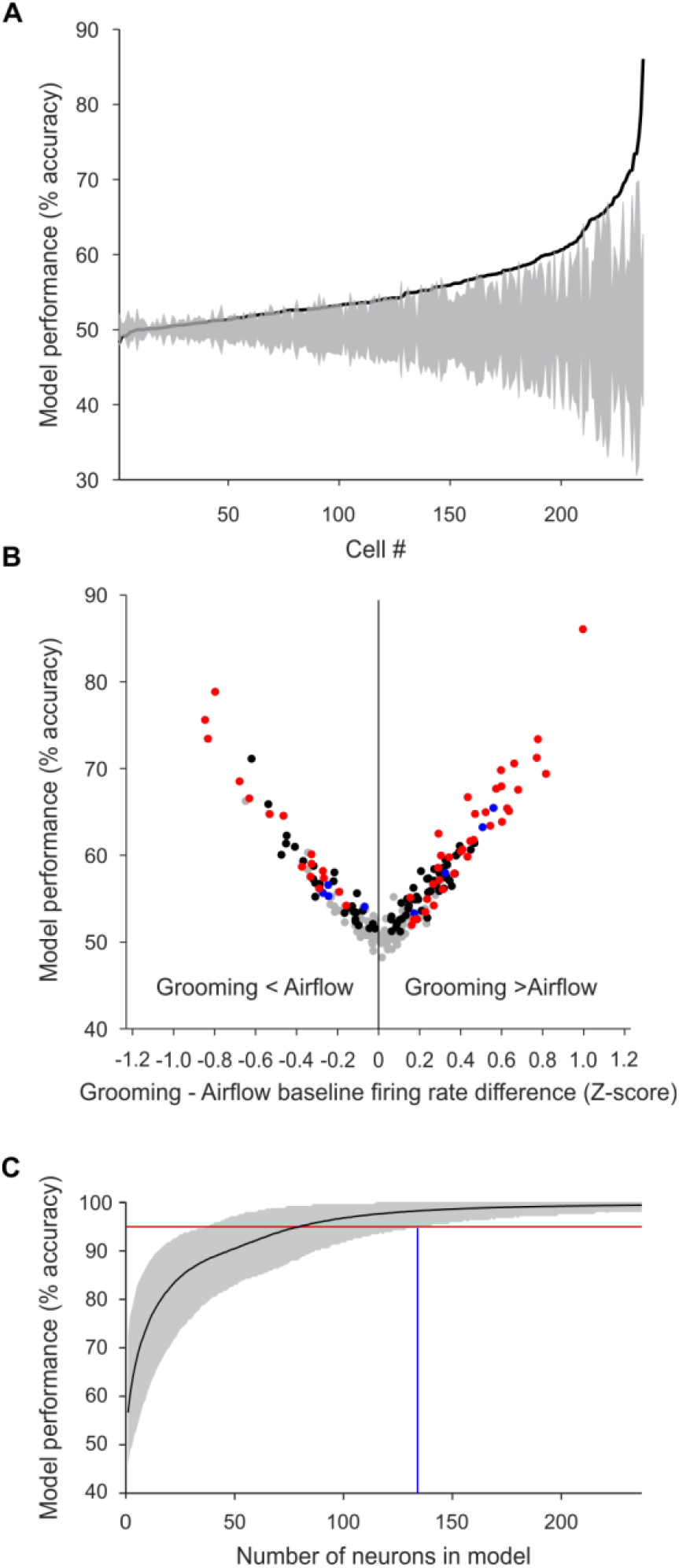
Context-related modulation of baseline-firing rates across the population of amygdala neurons. **A**. Performance of the SVM classifier for 237 individual neurons, sorted by decoding accuracy (black line). Gray shading corresponds to the 95% confidence bounds of the null distribution (note that the null distribution was generated individually for each cell). **B**. Performance of the SVM classifier for each of the neurons plotted as a function of the difference in baseline firing rates (normalized as Z-scores) between grooming and airflow blocks. The red dots (n = 52) represent cells that showed significant SVM accuracy and fulfilled the effect-size criteria for context-related activity. Black dots (n = 75) indicate cells that had significant SVM accuracy but did not pass the effect size criteria. Blue dots (n = 8) passed the effect size criteria but were not identified as having significant prediction accuracy based on the SVM. Gray dots (n = 102) showed no significant context-related activity when assessed using the SVM or the effect size criteria. **C**. The performance of the classifier on pseudo-populations of increasing numbers of neurons. The solid black line indicates the mean accuracy over 10,000 randomizations. Gray shading indicates the lower and upper bounds of the 95% confidence interval. The intersection of the red line (corresponding to 95% decoding accuracy) with the lower bound of the shading interval indicates the minimum number of neurons sufficient to accurately decode context at 95% (blue line, 134 neurons).

Figure 6B shows the performance of the SVM classifier for each of the 237 neurons plotted as a function of the difference in baseline firing rates (normalized as Z-scores) between grooming and airflow blocks. The correlation between these variables was strong for cases where the difference in baseline firing rates was negative (Pearson correlation rho = -0.96, p < 0.001) and where the difference was positive (Pearson correlation rho = 0.96, p < 0.001).

Furthermore, there was considerable overlap in neurons identified as exhibiting context-related baseline activity through SVM and those identified using the effect size criteria described above (Fig. 6B, red dots, n = 52 cells). An additional 75 cells (Fig. 6B, black dots) were identified has having significant SVM accuracy but did not pass the stringent effect size criteria. Only 8 cells passed the effect size criteria but were not identified as having significant prediction accuracy based on the SVM (Fig. 6B, blue dots). Finally, 102 cells (Fig. 6B, gray dots) showed no significant context-related activity when assessed using the SVM or using the effect size criteria.

Next, we determined the minimum number of randomly chosen neurons required for the accurate decoding of context. For increasing counts of neurons from 1 to 237, we generated 10,000 randomly chosen sets of baseline firing-rate values and determined the mean performance of the classifier (Fig. 6C). We found that a set of 134 neurons (blue vertical line in Fig. 6C) was necessary to yield correct classification of context above 95% accuracy (red horizontal line in Fig. 6C). The strong performance of the SVM was replicated in a principal component analysis (Fig. S3).

### Grooming and airflow stimuli elicit different autonomic states

We examined heart rate and heart rate spectral features in three monkeys to characterize their autonomic state during alternating blocks of grooming and airflow stimuli. Figure 7A shows an example recording of heart rate and the heart rate spectrogram over a 65-minute session. In this session, heart rate was lower during grooming (mean ± SD, 90 ± 10 BPM) than during airflow (101 ± 15 BPM, *t*-test *p* < 0.001). In addition, during grooming, we observed pronounced oscillations in the instantaneous heart rate around 0.3 Hz, a signature of respiratory sinus arrhythmia (RSA) linked to vagal tone or parasympathetic control of the heart (Fig 7A, lower panel) (Berntson et al., 2007). Parasympathetic states are typically associated with muscle relaxation, low vigilance, and social openness (Porges, 2001).

**Figure 7.**
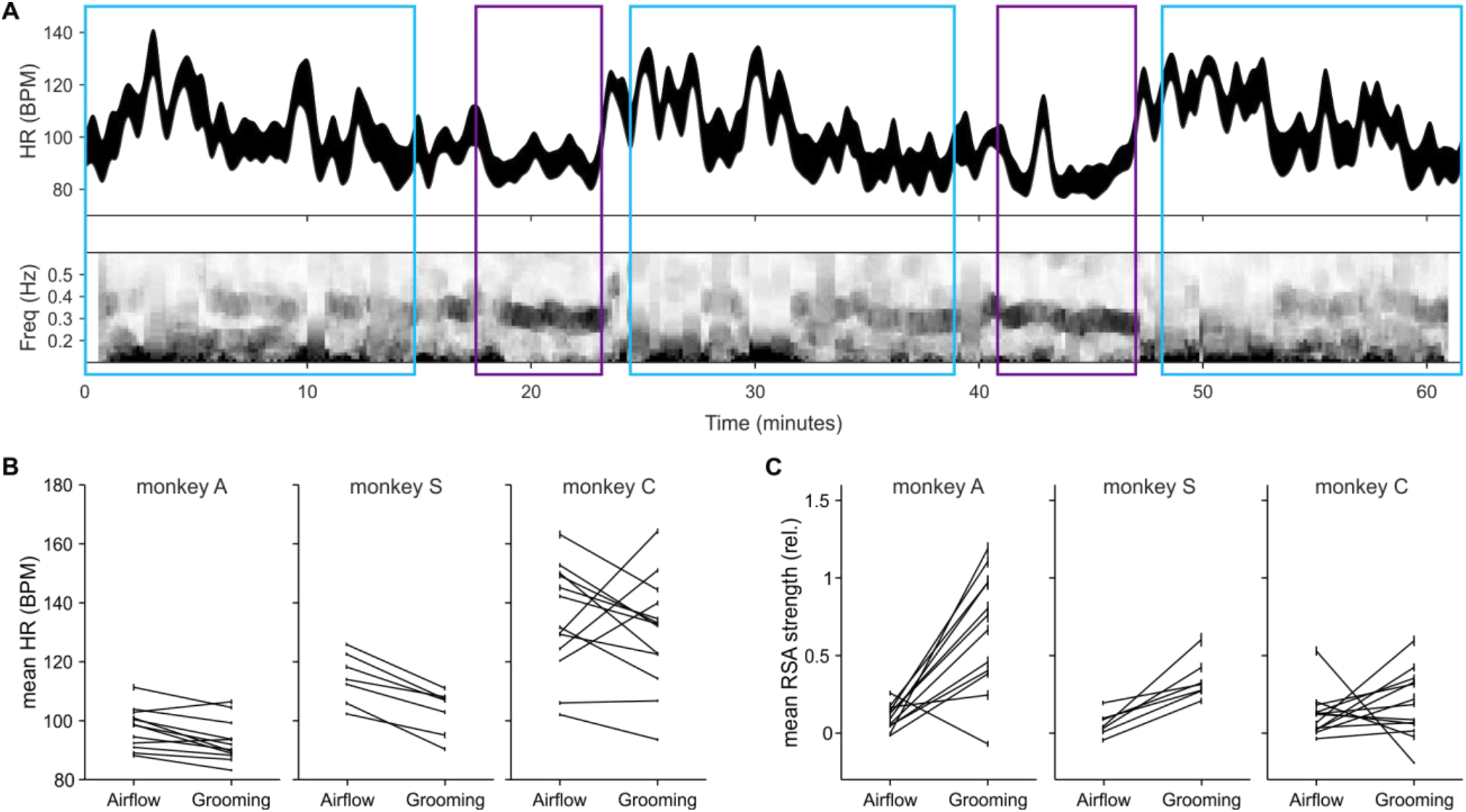
Autonomic states associated with airflow and grooming. **A**. Smoothed instantaneous heart rate ± sem from an example session (top), and heart rate variability spectrogram showing more pronounced respiratory sinus arrythmia (RSA, ∼ 0.3 Hz power) during grooming compared to airflow (bottom). Airflow and grooming blocks are indicated by blue and purple boxes, respectively. **B**. Mean heart rate in airflow and grooming blocks by session for monkeys A, S, and C (left to right). **C**. RSA strength relative to the mean across blocks, by session.

Across all sessions, average heart rate was lower in grooming blocks than in airflow blocks (monkey A: airflow 98 ± 7 BPM, grooming 92 ± 6 BPM, t-test p < 0.0001, n = 10 sessions; monkey S: airflow 114 ± 9 BPM, grooming 103 ± 8 BPM, p < 0.001, n = 7 sessions; monkey C: airflow 137 ± 20 BPM, grooming 124 ± 15 BPM, p < 0.001, n = 10 sessions). Exceptions to this pattern pertained to a few initial sessions when monkeys may not have been adjusted to the experimental situation (Fig. 7B; monkey A: first 2 sessions, monkey C: first 3 sessions; monkey S: did not show this pattern because he had been previously acclimated to electrophysiological recordings). Respiratory sinus arrhythmia, indicative of parasympathetic control of the heart, was also typically higher during grooming (Fig. 7C, all monkeys *p* < 0.05).

We considered whether absence of responses in the amygdala to grooming might be linked to the associated change in autonomic state during grooming. Although the autonomic state was different between bouts of grooming and airflow, moment-to-moment changes in the autonomic state were poor predictors of whether the neurons in the amygdala responded to grooming sweeps. For example, a brief increase in heart rate (Fig. 7A, minute 43 in the second grooming block) was not associated with reinstatement of responses to individual grooming sweeps. Across all sessions in the three monkeys for whom EKG was recorded, we identified episodes of high heart rate (see Methods) within the grooming blocks. A total of 31 such episodes were identified (episode duration 20.6 ± 12.3 s, range 10 – 50 s). Of the 37 amygdala neurons recorded during these episodes (all of which responded to airflow), none responded to grooming during these high heart rate periods.

## DISCUSSION

Here we offer a new perspective on sensory processing in the monkey amygdala brought about by experimental conditions that approximate grooming among primates. As grooming is an important affective and social stimulus for primates, and social-emotional stimuli elicit strong responses in the amygdala (e.g., Resnik and Paz; 2015; Gore et al., 2015; Chang et al., 2015; Kyriazi et al., 2018, Beyeler et al., 2018; Wang et al., 2017; Putnam and Gothard, 2019), we predicted that grooming would powerfully activate neurons in the amygdala. In addition to the social-emotional aspects of grooming, dynamic tactile sweeps across the skin were expected to evoke stronger responses than airflow because touch activates most non-nociceptive mechanoreceptive afferents whereas airflow primarily activates Pacinian corpuscles and hair follicle afferents (Vallbo et al. 1995; Delhaye et al. 2018). Contrary to our prediction, and despite autonomic signs of positive affect, neurons in the amygdala did not respond to individual grooming sweeps. The same neurons responded, however, to gentle airflow applied to the same area of the skin indicating that these neurons received tactile inputs from the face.

The mechanisms that account for the cessation of responses to grooming are unknown. Given that the fixed alternations of blocks and the sequencing of grooming stimuli were predictable, the suppression of responses might be attributed to the absence of a prediction error (e.g., Schultz et al., 1997, Averbeck and Costa, 2017). However, explicit tests for prediction-error encoding in the monkey amygdala showed that neurons sensitive to prediction errors continue to signal the identity of the associated stimulus (e.g., Belova et al., 2007; Tye and Janak, 2007). Furthermore, neurons in the amygdala, as well as dopaminergic neurons have been shown to respond to fully predictable rewards (e.g., Grabenhorst et al., 2012, Hamid et al., 2015). In our study, neurons recorded in control experiments (that violated the animal’s expectations and increased prediction error) remained unresponsive during grooming. Therefore, response modulation associated with prediction errors seems unlikely to account for the blunted responses during grooming.

A potent neuromodulator, like oxytocin, may be responsible for changing the neuronal responses between non-social and social touch (Handlin et al., 2021, Froemke & Young 2021). Indeed, the activity of oxytocinergic neurons increase during prosocial behaviors, including social touch (Hung et al. 2020, Tang et al. 2020). In the amygdala, oxytocin primarily increases the excitability of inhibitory interneurons (Hu et al. 2020; Crane et al. 2020) that could lead to suppression of principle cell activity in the amygdala during prosocial situations such as grooming, as observed in the present investigation.

Future experiments will be needed to explore the role of neuromodulators, control more substantially for the effects of predictability, and determine whether other, non-aversive stimuli can elicit a neural response during grooming blocks, when responses to individual grooming weeps were absent.

### The baseline activity of amygdala neurons may signal context

Overall, grooming induced two important changes in the activity of the amygdala. Neurons did not respond to individual touch stimuli through phasic departures from their baseline rate. Concurrently, in many neurons, baseline activity paralleled, through small but significant changes, the context in which the stimuli occurred. Although context in the present experiments likely had multiple components, the presence of the groomer alone was sufficient to elicit changes in baseline activity, regardless of the type of tactile stimulation received. This implies that baseline activity in the amygdala may be linked to the social context.

Classic field studies in primatology have demonstrated that social context can affect the meaning of sensory stimuli (Cheney and Seyfarth, 1990). Indeed, in different cognitive contexts, neurons in various brain areas respond differentially to the same visual stimuli (Rainer et al., 1998; Mante et al., 2013; Saez et al, 2015). Recently Jovanovic and colleagues (2022) showed that the baseline firing rates of prefrontal neurons in marmosets appear to signal social context. Our finding of persistent changes in baseline activity linked to social context is consistent with those findings. Furthermore, it reinforces previously demonstrated associations between long-lasting motivational states and modulation of baseline firing rates (Allen et al., 2019, Livneh et al., 2020). Anxiety, for example, is maintained over several seconds through the elevated baseline firing rate of neurons in the amygdala (Lee et al., 2017). Likewise, memory acquired through associative learning can persist for minutes through changes in baseline firing both in the amygdala and the anterior cingulate cortex (Taub et al, 2018). Similarly, baseline activities of neurons in the basal ganglia and the cortex retain information about the outcome from multiple preceding trials to guide future behavioral choices (Histed et al., 2019). In the insula, baseline activity tracks the satiety of animals (Livneh et al., 2020). Furthermore, baseline activity in multiple brain areas appears to encode the engagement of the animal with external stimuli (Steinmetz et al., 2019).

The cellular mechanisms that give rise to these persistent changes in baseline activity in the amygdala are unknown. Baseline firing rates depend on the level of activity in the local network within which neurons are embedded (e.g., Tsodyks et al., 1999) but also on inputs from larger networks that govern global brain states (John et al., 2016; Barrett and Satpute, 2013). Projections from the brainstem, hypothalamus, or from the nucleus basalis of Meynert (Amaral et al.,1992) that release neuromodulators into the amygdala (Hariri and Whalen; Turchi et al., 2018; Crouse et al., 2020) may be candidates for cellular processes that govern baseline activity.

### Social touch is associated with positive affective state

By virtue of its role as the major hub in the brain where multiple networks intersect (Bickart et al., 2014), the amygdala has nuanced control over vigilance (Davis and Whalen, 2001), emotion (LeDoux, 1992), and social behavior (Adolphs, 2001; Dunbar 2001; Gothard, 2020). Although our subjects were seated in a primate chair and were groomed by humans, their physiological responses resembled that of macaques groomed by conspecifics in natural settings. They showed vagal tone, slowed heart rate, and muscular relaxation (Boccia, 1989; Schino and Aureli, 2007; Grandi and Ishida, 2015). In natural settings, the recipient of grooming typically relinquishes attentive scanning of the environment to the groomer (Boccia, 1989). In contrast, during vigilant scanning of the environment, the amygdala seems to prepare the organism to detect and respond to salient, behaviorally meaningful stimuli (Davis and Whalen, 2001).

The lack of vigilance and the presence of vagal tone might partially account for the cessation of neuronal responses to individual tactile stimuli during grooming. Under these conditions, naturalistic grooming touches might be processed by the amygdala not as individual tactile stimuli but as a continuous stimulus that elicits a prolonged internal state. Another possibility is that while grooming-related touches were suppressed, unpredicted touches to other body regions might not be. Such behaviorally dependent gating of cutaneous input has been observed in cuneate nucleus, a site containing second order somatosensory neurons in the lemniscal pathway (He et al., 2022). In the amygdala, suppression of tactile sensitivity may reflect a switch from sensing individual stimuli to ‘feeling’ an internal state. Future studies, on the mechanism of sensory suppression or gating hold promise for explaining the phenomenon reported here.

## ACKNOWLEDGEMENTS

Supported by 1R56MH115681 and R01MH121009

We thank Natalia Noel Jacobson for spike sorting. Gowri Somasekhar helped record data from monkey C. Seunghyun Lee helped set up and troubleshoot the equipment. Derek O’Neill helped with surgeries, MRI’s, and general technical support. We consulted Dr. Ueli Rutishauser on optimal ways to apply SVM to our data and Drs. Francesco Battaglia and Bryan Souza for verifying the outcome of the machine learning algorithm.

## AUTHOR CONTRIBUTIONS

ABM: Collected a subset of data, curated and pre-processed part of the data, performed all final analyses, made all the figures, supervised the work of MC and AIB, wrote the manuscript

MAC: Collected a subset of data, performed part of the analysis, curated and pre-processed part of the data, helped prepare the manuscript

RKA: Designed the experimental protocol, collected a large portion of the data, curated and pre-processed part of the data, performed anatomical reconstruction of recording sites

AIB: Collected a subset of data, curated and pre-processed a subset of the data, performed part of the analysis, helped prepare the manuscript

EAH: Trained the animals, collected a large portion of the data, curated and pre-processed part of the data

SB: Helped secure funding, provided expertise and feedback, reviewed and edited multiple versions of the manuscript

AJF: Conceptualization, helped secure funding, provided expertise and feedback, manufactured and tested custom hardware, wrote, reviewed, and edited multiple versions of the manuscript

KMG: Conceptualization, secured funding, supervised data acquisition, pre-processing, analysis, and visualization, wrote, reviewed, and edited multiple versions of the manuscript

## DECLARTION OF INTERESTS

None of the authors have conflicting interests

## STAR METHODS

## EXPERIMENTAL MODEL AND SUBJECT DETAILS

### Subjects and Surgical procedures

Four adult male rhesus macaques, E, A, C, and S (weights 13.7, 13.6, 10.6, and 12.6 kg; ages 8, 12, 6, and 5 years respectively), were prepared for neurophysiological recordings from the amygdala and somatosensory cortex. The stereotaxic coordinates of the right somatosensory cortex and right amygdala in each animal were determined (left and right amygdala in monkey C) based on 3T structural magnetic resonance imaging (MRI) scans (isotropic voxel size = 0.5 mm) (Fig. S4, S5). A square (26 × 26 mm inner dimensions) polyether ether ketone (PEEK) MRI compatible recording chamber was surgically attached to the skull and a craniotomy was made within the chamber. The implant also included three titanium posts, used to attach a ring for head fixation to the implant. Between recording sessions, the craniotomy was sealed with a silicone elastomer (Kwick-Sil, WPI) to prevent growth and scarring of the dura (Spitler and Gothard, 2008). All procedures complied with NIH guidelines for the use of non-human primates in research and were approved by the University of Arizona’s Institutional Animal Care and Use Committee.

## METHOD DETAILS

### Electrophysiological procedures

Single-unit spiking activity was recorded using linear electrode arrays (V-probes, Plexon Inc, Dallas, TX) with 16 (16 sessions) or 32 (32 sessions) equidistant contacts at 400 μm (16 contacts) or 200 μm (32 contacts) separation along a 260 μm diameter shaft. Data were collected using Plexon OmniPlex data acquisition hardware and software (RRID:SCR_014803). Electrode arrays were acutely lowered into the right somatosensory cortex and amygdala for each recording session using a Thomas Recording Motorized Electrode Manipulator (Thomas Recording GmbH, Giessen, Germany). Impedance for each contact ranged from 0.2 to 1.2 MΩ. The anatomical location of each electrode was calculated by aligning a scaled image of the chamber to a series of coronal MR images and fiducial markers (co-axial columns of high contrast material). During recordings, slip-fitting grids with 1 mm distance between cannula guide holes were placed in the chamber, allowing sampling of medio-lateral and anterior-posterior locations in the amygdala and somatosensory cortex. A twenty-three-gauge cannula was inserted through the guide holes and advanced 4-6 mm below the dura into the cortex. V-probes were then driven through the cannula and to the amygdala or somatosensory cortex at a rate of 70-100 µm/s, slowing to 5-30 µm/s after the tip of the V-probe crossed into the estimated location of the central nucleus of the amygdala or primary somatosensory cortex. Data were recorded in 48 sessions: monkey E = 9 (amygdala and somatosensory cortex), A = 13 (amygdala and somatosensory cortex), C = 13 (left and right amygdala), S = 7 (2 probes in right amygdala) and 6 (2 probes in right somatosensory cortex). The analog signals from each channel on the V-probe were digitized at the headstage (Plexon Inc, HST/16D Gen2) before being sent through a Plexon pre-amplifier, filtering from 0.3 - 6 kHz and sampling continuously at 40 kHz. The raw data derived from these recordings are available upon request. Single units were sorted offline (Plexon offline sorter version 3, RRID:SCR_000012) predominately using principal component analysis, and spike times were rounded to the nearest millisecond.

### Autonomic recordings

In three monkeys (A, C, and S), heart rate was recorded using self-adhesive H59P electrocardiogram (ECG) electrodes (Cardinal Health, Waukegan, IL) attached to two shaved skin patches on the animal’s back and recorded at 1 kHz. R waves, corresponding to ventricular contraction, were manually discriminated using the Plexon offline sorter.

### Stimulus delivery

Monkeys were seated in a primate chair and placed in a recording booth featuring a custom-made airflow delivery apparatus (Crist Instruments), as described in detail previously (Morrow et al., 2019). The airflow system delivered gentle, non-aversive airflow stimuli via computer-controlled solenoid valves through low pressure vinyl tubing that was fed through Loc-line hoses (Lockwood Products). The Loc-line nozzles were placed ∼2 cm from the monkeys’ fur, aimed at ten locations on the face and head, avoiding the eyes, ears, and nose. Nozzles were placed to the left and right, adjacent to the lower muzzle, upper muzzle, brow, lateral head, and posterior head. An additional nozzle was placed behind the monkey to act as a sham control for alerting and auditory responses. The pressure detected 2 cm away from the nozzle was about 10 Pa (roughly equivalent to the pressure delivered by a gentle breeze traveling at 3 m/s). Delivery of airflow stimuli was controlled using custom-written code in Presentation Software (Neurobehavioral Systems). The airflow stimuli were delivered for 1s (in most sessions, except for some early sessions when airflow lasted for 1.5s) followed by a 3 second inter-stimulus interval. Airflow stimuli were pseudo-randomly delivered to each of the 11 locations (including sham) as shown in Figure 1. Juice reward was delivered at the end of each set. Each set of 11 stimuli was repeated 10 times, for one airflow block.

For grooming blocks, the monkey’s trainer entered the recording booth, sat in front of the animal, and delivered gentle, grooming-like sweeps to the monkey’s face. Grooming sweeps lasted 1 – 2 s, followed by a 2 – 4 s inter-stimulus interval. Typically, the monkey received 10 grooming sweeps to the left upper muzzle, followed by 10 sweeps to the left brow. (In monkey C, right upper muzzle and brow were included as we recorded from the amygdala in both hemispheres). The main reason for not applying the grooming stimuli to all face regions was the difficulty (and excessive time) in removing and re-positioning the air nozzles around the monkey’s face to give access to the experimenter’s hand for grooming. Each set of sweeps was repeated 5 times in one block. The timing and the contact pressure of the grooming sweeps were recorded using a custom-built pressure-transducer placed on the pad of the index finger inside a vinyl glove. The contact pressure of the grooming sweeps gradually increased and then decreased (see Fig. 3A), with a peak pressure between 0.5-2 kPa. The onset of the rise in contact pressure from the pressure transducer on the groomer’s finger was used as time zero for the alignment of spiking activity to the grooming stimulus. The groomer attempted to mimic the grooming gesture of monkeys in terms of sweep speed, duration, and consistent delivery of the grooming sweep to the same skin location. As expected from manual application, there was some variability in pressure and sweep duration across trials. At the end of the grooming block the monkey received food reward (e.g., 2 - 3 peanuts) and the trainer exited the recording booth.

In a typical experimental session, three airflow blocks were interspersed with two grooming blocks. Typically, the firs block was an airflow block. The pattern was varied in about 1/3 of sessions (for example, the session might start with a grooming rather than an airflow block). The durations of airflow and grooming blocks were also varied in some sessions. In a subset of sessions, control blocks were appended to the end of a standard set of blocks. To evaluate the effects of contact pressure on neural responses, we decreased or increased the airflow pressure to half and double the standard pressure. Similarly, we decreased or increased the pressure of the grooming sweeps to approximately half or double the usual pressure. Pressure controls were performed in three sessions of monkey A. To control for visual inputs from the looming hand of the groomer, we blindfolded the monkey during a set of airflow-grooming-airflow blocks (monkey E: 2 sessions, monkey S: 2 sessions, monkey A: 2 sessions). To control for the effect of differing reward contingencies between airflow and grooming, the reward schedule was reversed: instead of receiving reward between each set of airflow stimuli, the monkey received reward at the end of the entire block. Likewise, instead of receiving reward at the end of the grooming block, the monkey received rewards between each set of 10 grooming sweeps (monkey C: 2 sessions). An additional control was designed where airflow stimuli were delivered in the same pattern as standard grooming stimuli (10 stimuli to the left upper muzzle followed by 10 stimuli to the left brow, repeated 5 times), and the grooming stimuli were delivered pseudo-randomly to 8 locations, repeated 10 times (monkey A: 1 session, monkey C: 1 session, monkey S: 3 sessions).

All monkeys were trained for several weeks prior to electrophysiological recordings to acclimate them to the airflow puffers, grooming, and EKG electrodes.

## QUANTIFICATION AND STATISTICAL ANALYSIS

### Quantification of neural responses

We recorded 615 neurons from the amygdala (monkeys E: 58, A: 148, C: 234, S: 175) and 375 neurons from the somatosensory cortex (monkeys E: 63, A: 161, S: 151). A neuron was included in the analysis if it met the following criteria:

1. minimum firing rate: the average firing rate across the experiment was at least 1 Hz
2. criterion for responsivity to tactile stimuli: stable for at least 10 trials of each airflow and grooming stimulus locations.

Spike times and waveforms were imported into MATLAB for analysis using scripts from the Plexon-MATLAB Offline Software Development Kit (Plexon). All analyses were conducted using custom scripts in MATLAB R2021b (Mathworks). Colors were tested for color-blind friendliness using online algorithms at *Coloring for Colorblindness* (https://davidmathlogic.com/colorblind).

### Grooming and airflow responses

Stimulus induced responses were identified by comparing pre-stimulus and post-stimulus firing rates. The pre-stimulus window was defined as 1000 ms to 250 ms before stimulus onset; the post-stimulus window was defined as 200 ms after stimulus onset to the end of the stimulus. The average firing rate of each of these windows was compared using a paired *t*-test. A neuron was classified as having a stimulus induced response if, in at least one stimulus location, the stimulus-baseline comparison was significant at *p* < 0.05 and the mean stimulus rate was at least 1 Hz different from the mean baseline rate.

To compare responses between grooming and airflow stimuli, we considered airflow responses only at locations that were also grooming locations. Although we delivered airflow stimuli to 10 locations, and required at least 10 stimulations of each location, we included in the analysis responses only to the airflow locations that were also groomed e.g., the left upper muzzle and left brow.

We compared responses during control and standard experimental conditions using only neurons that were active in both conditions. Proportions of cells that responded were compared using the chi-squared test of proportions.

### Baseline activity

The criterion for including neurons in baseline firing rate analysis was stable firing for 4 blocks, (two airflow and two grooming blocks in alternations). Concatenated segments of baseline firing rates for two grooming and two airflow blocks were generated by removing stimulus windows (with a 300-ms buffer before and after), inter-block intervals, and reward windows. The resulting baseline firing rates were binned in 1 s bins and converted to Z-scores.

### Context-related baseline effects

In the analysis of context-related modulation, we included 237 neurons from the amygdala that were stable for 2 airflow and 2 grooming blocks (monkey A, n = 56; monkey C, n =78; monkey S, n = 103). We included neurons with at least 60 baseline bins (1 s bins) in each of the four blocks.

We calculated context-related effect sizes as Cohen’s *d* for independent samples (*d*_*s*_) (as in Lakens, 2013):

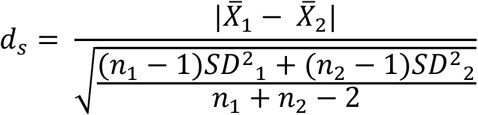

where 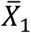 and 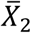 are the mean baseline firing rates in the two sets of interest. For each cell, we computed three values of *d*_*s*_: 1) comparing grooming to airflow to identify the size of context-related effects, 2) comparing the first grooming block to the second grooming block to identify the size of between-grooming-block variability, and 3) comparing the first airflow to the second airflow block to identify the size of between-airflow-block variability. We classified cells as exhibiting context-related responses if the grooming-to-airflow effect size was greater than the standard minimum threshold of 0.2 (Cohen, 1992) and the grooming-to-airflow effect size was at least 1.5 times greater than the largest effect size of either the grooming-to-grooming or airflow-to-airflow block comparisons (Fig. S2).

### Support vector machine classifier (SVM) and principal component analysis (PCA) to characterize context responses

To determine whether context could be decoded from the baseline firing rates of amygdala neurons, we trained a support vector machine (linear classifier). We used a 10-fold cross-validation on balanced counts of airflow and grooming bins for each cell. For each neuron we also generated a null distribution by randomly assigning bins to either grooming or airflow blocks, repeated 10,000 times, and compared the upper bound of the resulting 95% confidence interval to the performance when using correctly assigned bins (note that the 10-fold cross-validation can result in performance below 50%, however the null distribution was centered on 50% for all neurons).

To identify the minimum population size required for accurate decoding of context, we determined the SVM performance using sets of 1 to 237 neurons. Balanced counts of 60 bins per block were used for all cells. For each set size, we generated 10,000 sets of randomly selected neurons from our recorded population of 237 (with replacement) and found the mean performance of the classifier and 95% confidence interval of the bootstrapped distribution. (We repeated the analysis without replacement and found no qualitative difference in overall results.) We used the lower bound of the 95% confidence interval to determine the performance of the classifier for that set size.

We applied principal component analysis (PCA) to visualize the separation of baseline activity in airflow and grooming bins (Fig. S3). The separability of the resulting projections of the population activity onto the first component was assessed using k-means clustering. The accuracy of the clustering was determined by comparing the result of the k-means clustering to the veridical bin assignments to airflow or grooming.

### Heart rate and respiratory sinus arrhythmia (RSA)

Instantaneous heart rate was calculated as the inverse of the duration of each inter-beat interval (IBIs). Noise and movement artifacts were removed, as well as windows with biologically implausible IBIs (greater than 1500 ms and less than 250 ms, i.e., instantaneous heart rates less than 40 beats per minute (BPM) or greater than 240 BPM). Instantaneous heart rates were interpolated to 1 ms timescales using a modified Akima cubic Hermite interpolation. Mean heart rates between conditions were compared using a paired *t*-test across sessions.

Heart rate variability was calculated from noise-free heartbeat times in sliding windows of 60 s with a 3 s step, using a multi-taper power spectral density estimate with 7 Slepian tapers for ± 0.07 Hz smoothing. For each spectrum, we identified peaks between 0.25 and 0.5 Hz, corresponding to respiratory rates between 15 and 30 breaths per minute. RSA strength in that time window was defined as the mean power at the peak ± half-width. The RSA strength at each time was normalized to the median strength across all time steps: *RSA*(*t*)=(*RSA strength* (*t*) − *μ*)/*μ*.

### Episodes of high heart rate during grooming

We identified episodes of high heart rate during grooming using the following approach. We first determined the mean (and SD) heart rate during prolonged periods of stable heart rate (mean ± SD = 11.4 ± 7.0 minutes, range = 6 - 20 minutes) within the grooming blocks. All grooming periods within a session were then scanned for epochs for which heart rate was > 2 SD above the stable-period mean and with a duration of at least 10 s. This process identified a total of 31 high-heart rate episodes in the three monkeys for whom EKG was recorded. Peri-stimulus time histograms of neural activity were generated for grooming sweeps during these epochs of high heart rate. We then applied the same criteria as described above (“Grooming and airflow responses”) to identify whether neurons responded to grooming during these epochs.

## SUPPLEMENTAL INFORMATION

### 1. Control experiments

**Figure S1.**
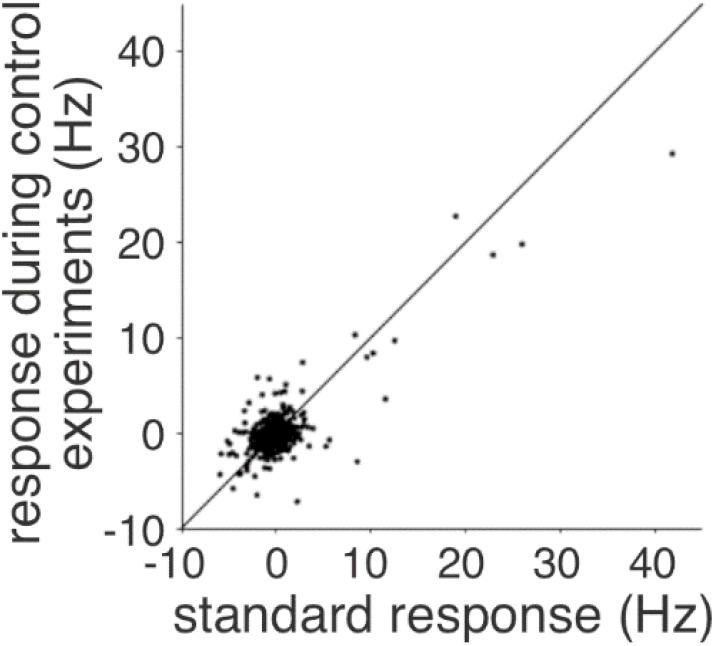
Firing rate responses for 188 amygdala neurons recorded during control experiments plotted as a function of the responses during standard conditions (see examples in Fig 3). Control experiments involved changes in grooming pressure (n = 18), blindfolding (n = 40), changing reward contingencies (n = 40), changing predictability of stimuli (n = 36), and having the groomer sit in the booth during airflow blocks (n = 54). The responses during the standard experiment were highly correlated with the responses during the controls (Pearson correlation rho = 0.78, p < 0.0001). Therefore, manipulating various aspects of the experiment protocol did not influence the responses of the recorded neurons.

### 2. Identification of context-related changes in baseline firing rate

For a neuron to be considered as exhibiting context-related activity in baseline firing, the grooming-to-airflow effect size in baseline firing rates needed to exceeded 0.2 (Cohen, 1992) and be 1.5 times greater than the largest of the within condition measures of effect size. Fig. S2 shows the effect size for grooming-to-airflow conditions for each amygdala neuron plotted as a function of the largest effect size of either the grooming-to-grooming or airflow-to-airflow block comparisons (X-X, Fig. S2). The dashed horizontal line indicates the minimal effect-size threshold for grooming – airflow block comparisons (*d*_*s*_ = 0.2). The diagonal line represents 1.5 times the X-X effect size. Red dots indicate cells (n = 60) that fulfilled both criteria and, therefore, were considered as exhibiting context-related activity in baseline firing rates.

**Figure S2.**
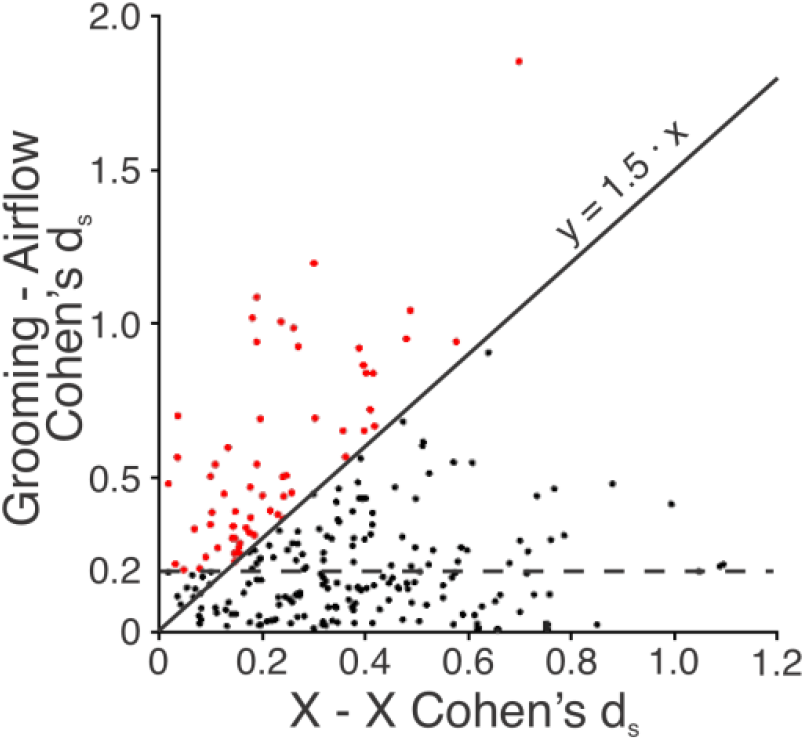
Effect size for grooming-to-airflow comparison in baseline firing rates for 237 amygdala neurons plotted as a function of the largest effect size of either the grooming-to-grooming or airflow-to-airflow block comparisons (X-X). Dashed horizontal line represents the minimum effect size for grooming – airflow comparisons. Diagonal line represents 1.5 times the X-X effect size. To be considered possessing context-related activity in baseline firing, neurons had to exceed the effect size threshold and to have an effect size for grooming – airflow at least 1.5 times larger than the X-X effect size (red dots, n = 60).

### 3. Context can be decoded from baseline firing rates despite minimal explained variance

Given that the SVM decoded context from the baseline firing rates of the population of 237 amygdala neurons, we asked how explanatory is context of the variance in the baseline firing rate? We applied principal component analysis (PCA) to the same bins as in the SVM and applied k-means clustering to the first component. The first principal component of PCA could be correctly classified using k-means clustering at 95.4% accuracy. As shown in Figure S2, the projections of the first two components were separable into grooming and airflow bins (purple and blue dots, respectively) on the axis corresponding to the first component (the x-axis). The decision boundary of k-means clustering is indicated by the black line at 0 on the first component. The first component explained 8% of the overall baseline firing rate variance, corresponding to an average of 0.5 Hz difference in baseline firing rates between grooming and airflow blocks. Thus, even small variations in baseline activity carry useful information about context for downstream structures. Of the 60 cells identified as having context-related activity in baseline activity (see Fig. S2), the mean (± SD) difference in baseline firing rates was 1.2 ± 1.2 Hz.

**Figure S3.**
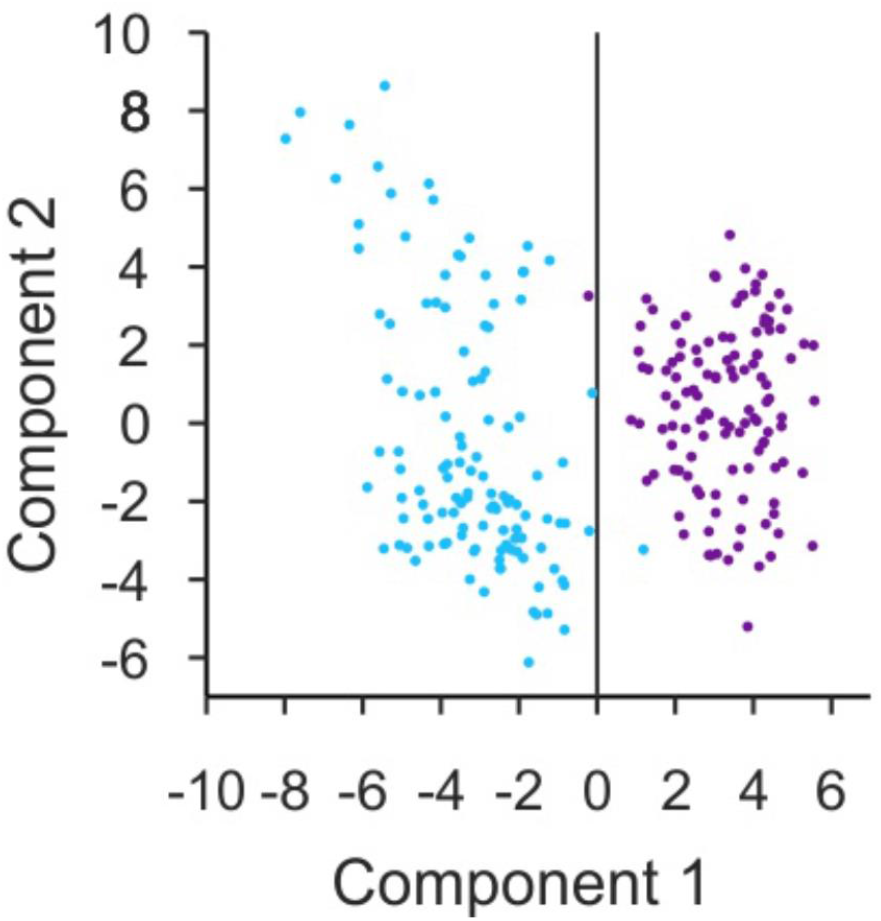
Principal component and k-means clustering of population baseline firing rates from grooming (purple) and airflow (blue) time bins. K-means clustering of the first component values into grooming or airflow time bins was accurate for 95.4% of bins, indicated by the clustering boundary at 0 (black line).

### 4. Reconstruction of recording sites

From the structural MRI scans (isotropic voxel size of 0.5mm), an atlas was built for each monkey. The atlas contained all MRI slices that contained the amygdala or area 3b of the primary somatosensory cortex. The boundaries of the main nuclei of the amygdala were outlined on each MRI slice (Fig. S4). Anatomical reconstructions of electrode targets were based on post-surgical MRIs that used columns of contrast positioned coaxially with the recording chambers, allowing us to calculate the x-y-z location of each recording site in the amygdala relative to the chamber coordinates (error magnitude maximum 1 mm). The reconstructed locations of the electrode contacts that recoded the amygdala neurons included in this study are shown in figure S4 and those for SI in figure S5.

**Figure S4.**
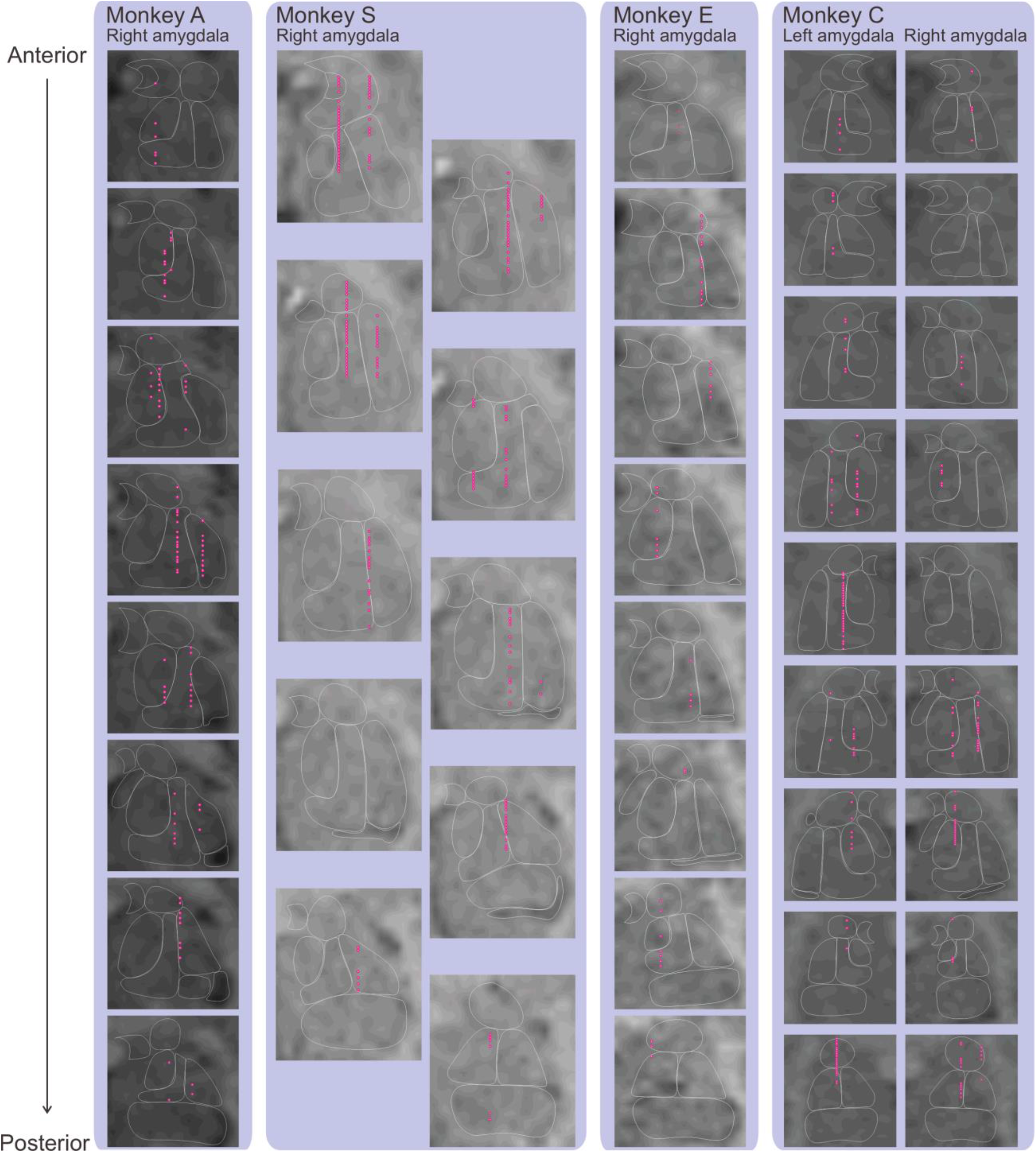
Anatomical reconstruction of amygdala recording sites for four monkeys. MRI slices are shown from most anterior (top) to most posterior (bottom). White lines indicate the estimated boundary of amygdala nuclei. Red dots indicate the estimated locations of electrode contacts that recorded neurons included in this study.

**Figure S5.**
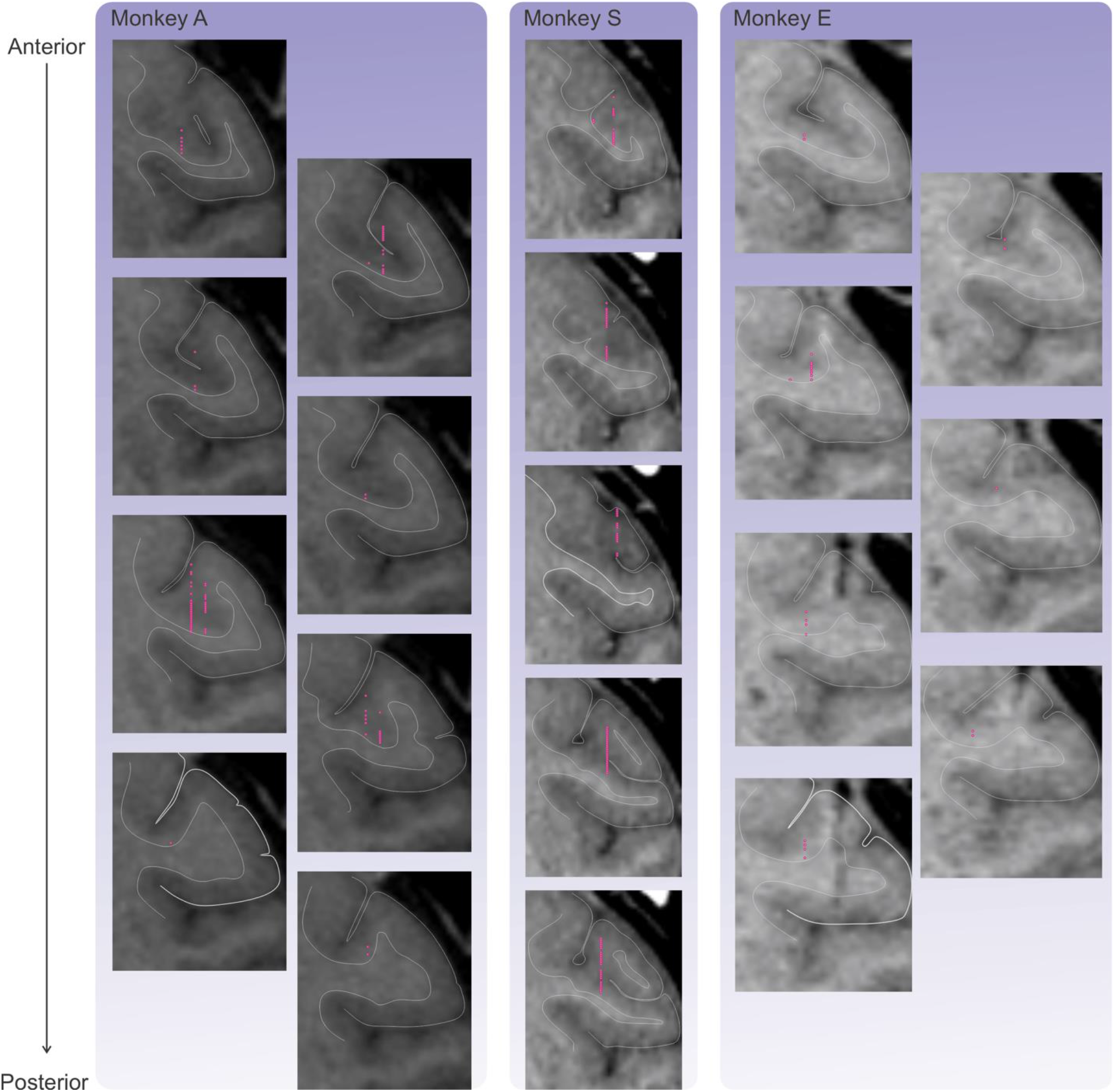
Anatomical reconstruction of SI recording sites for three monkeys. MRI slices are shown from most anterior (top) to most posterior (bottom). Red dots indicate the estimated locations of electrode contacts that recorded neurons included in this study.

## REFERENCES

Adolphs, R. (2001). The neurobiology of social cognition. Current opinion in neurobiology,11(2), 231–239. 10.1016/s0959-4388(00)00202-6

Allen, W. E., Chen, M. Z., Pichamoorthy, N., Tien, R. H., Pachitariu, M., Luo, L. & Deisseroth, K. (2019). Thirst regulates motivated behavior through modulation of brainwide neural population dynamics. Science, 364(6437), 0–10. https://doi.org.10.1126/science.aav3932

Amaral DG, Price JL, Pitkanen A, Carmichael TS: Anatomical organization of the primate amygdaloid complex. In The Amygdala: Neurobiological Aspects of Emotion, Memory, and Mental Dysfunction. Edited by Aggleton JP. Wiley-Liss; 1992.

Amir, A., Headley, D. B., Lee, S. C., Haufler, D., & Paré, D. (2018). Vigilance-Associated Gamma Oscillations Coordinate the Ensemble Activity of Basolateral Amygdala Neurons. Neuron,97(3), 656-669.e7. https://doi.org/10.1016/j.neuron.2017.12.035

Aureli, F., Preston, S. D., & de Waal, F. B. M. (1999). Heart Rate Responses to Social Interactions in Free-Moving Rhesus Macaques (Macaca mulatto): A Pilot Study. In Journal of Comparative Psychology (Vol. 113, Issue 1). 10.1037/0735-7036.113.1.59

Averbeck, B. B., & Costa, V. D. (2017). Motivational neural circuits underlying reinforcement learning. Nature Neuroscience,20(4), 505–512. https://doi.org/10.1038/nn.4506

Bales, K. L., Witczak, L. R., Simmons, T. C., Savidge, L. E., Rothwell, E. S., Rogers, F. D., Manning, R. A., Heise, M. J., Englund, M., & Arias del Razo, R. (2018). Social touch during development: Long-term effects on brain and behavior. In Neuroscience and Biobehavioral Reviews (Vol. 95, pp. 202–219). Elsevier Ltd. https://doi.org/10.1016/j.neubiorev.2018.09.019

Barrett, L. F., & Satpute, A. B. (2013). Large-scale brain networks in affective and social neuroscience: Towards an integrative functional architecture of the brain. In Current Opinion in Neurobiology (Vol. 23, Issue 3, pp. 361–372). https://doi.org/10.1016/j.conb.2012.12.012

Belova, M. A., Paton, J. J., Morrison, S. E., & Salzman, C. D. (2007). Expectation modulates neural responses to pleasant and aversive stimuli in primate amygdala. Neuron,55(6), 970–984. https://doi.org/10.1016/j.neuron.2007.08.004

Berntson, G. G., Cacioppo, J. T., & Grossman, P. (2007). Whither vagal tone. Biological Psychology,74(2), 295–300. https://doi.org/10.1016/j.biopsycho.2006.08.006

Beyeler, A., Chang, C. J., Silvestre, M., Lévêque, C., Namburi, P., Wildes, C. P., & Tye, K. M. (2018). Organization of Valence-Encoding and Projection-Defined Neurons in the Basolateral Amygdala. Cell Reports,22(4), 905–918. https://doi.org/10.1016/j.celrep.2017.12.097

Bickart, K. C., Dickerson, B. C., & Barrett, L. F. (2014). The amygdala as a hub in brain networks that support social life. In Neuropsychologia (Vol. 63, pp. 235–248). Elsevier Ltd. https://doi.org/10.1016/j.neuropsychologia.2014.08.013

Boccia, M. L., Reite, M., & Laudenslager, M. (1988). On the Physiology of Grooming in a Pigtail Macaque. In Physiology & Behavior (Vol. 45). 10.1016/0031-9384(89)90089-9

Callaghan, B. L., & Tottenham, N. (2016). The Stress Acceleration Hypothesis: Effects of early-life adversity on emotion circuits and behavior. In Current Opinion in Behavioral Sciences (Vol. 7, pp. 76–81). Elsevier Ltd. https://doi.org/10.1016/j.cobeha.2015.11.018

Caldji, C., Tannenbaum, B., Sharma, S., Francis, D., Plotsky, P. M., & Meaney, M. J. (1998). Maternal care during infancy regulates the development of neural systems mediating the expression of fearfulness in the rat. Proceedings of the National Academy of Sciences,95(9), 5335–5340. https://doi.org/10.1073/pnas.95.9.5335

Cascio, C. J., Moore, D., & McGlone, F. (2019). Social touch and human development. Developmental Cognitive Neuroscience, 35, 5–11. https://doi.org/10.1016/j.dcn.2018.04.009

Chang, S. W. C., Fagan, N. A., Toda, K., Utevsky, A. v., Pearson, J. M., Platt, M. L., & Gazzaniga, M. S. (2015). Neural mechanisms of social decision-making in the primate amygdala. Proceedings of the National Academy of Sciences of the United States of America,112(52), 16012–16017. https://doi.org/10.1073/pnas.1514761112

Cheney, D. L., & Seyfarth, R. M. (1990). How monkeys see the world: Inside the mind of another species. University of Chicago Press.

Cohen, J. A power primer (1992). Psychological Bulletin, 112, 155–159.

Crane JW, Holmes NM, Fam J, Westbrook RF & Delaney AJ (2020). Oxytocin increases inhibitory synaptic transmission and blocks development of long-term potentiation in the lateral amygdala. J Neurophysiol 123: 587–599.

Crouse, R. B., Kim, K., Batchelor, H. M., Girardi, E. M., Kamaletdinova, R., Chan, J., Rajebhosale, P., Pittenger, S. T., Role, L. W., Tamage, D. A., Jing, M., Li, Y., Gao, X. B., Mineur, Y. S. & Picciotto, M. R. (2020) Acetylcholine is released in the basolateral amygdala in response to predictors of reward and enhances the learning of cue-reward contingency. eLife, 9, 1–31 https://doi.org/10.7554/ELIFE.57335

Dagnino-Subiabre, A. (2022). Resilience to stress and social touch. In Current Opinion in Behavioral Sciences (Vol. 43, pp. 75–79). Elsevier Ltd. https://doi.org/10.1016/j.cobeha.2021.08.011

Davis, M., & Whalen, P. J. (2001). The amygdala: vigilance and emotion. In Molecular Psychiatry (Vol. 6).

Delhaye, B.P., Long, K.H., and Bensmaia, S.J. (2018). Neural Basis of Touch and Proprioception in Primate Cortex. Compr Physiol 8, 1575–1602. 10.1002/cphy.c170033.

Dunbar, R. I. M. (2010). The social role of touch in humans and primates: Behavioural function and neurobiological mechanisms. In Neuroscience and Biobehavioral Reviews (Vol. 34, Issue 2, pp. 260–268). https://doi.org/10.1016/j.neubiorev.2008.07.001

Dunbar, R. I. M. (2020). Structure and function in human and primate social networks: implications for diffusion, network stability and health. Proceedings of the Royal Society A: Mathematical, Physical and Engineering Sciences,476(2240), 20200446. https://doi.org/10.1098/rspa.2020.0446

Froemke RC & Young LJ (2021). Oxytocin, Neural Plasticity, and Social Behavior. Annu Rev Neurosci 44: 1–23.

Gangopadhyay, P., Chawla, M., Dal Monte, O., & Chang, S. W. C. (2021). Prefrontal–amygdala circuits in social decision-making. In Nature Neuroscience (Vol. 24, Issue 1, pp. 5–18). Nature Research. https://doi.org/10.1038/s41593-020-00738-9

Gee, D. G. (2016). Sensitive Periods of Emotion Regulation: Influences of Parental Care on Frontoamygdala Circuitry and Plasticity. In New Directions for Child and Adolescent Development (Vol. 2016, Issue 153, pp. 87–110). John Wiley and Sons Inc. https://doi.org/10.1002/cad.20166

Gordon, I., Voos, A., Bennett, R., Bolling, D., Pelphrey, K., Kaiser, M. (2013), Brain mechanisms for processing affective touch. Human Brain Mapping, 34, 914–922. DOI: 10.1002/hbm.21480

Gore, F., Schwartz, E. C., Brangers, B. C., Aladi, S., Stujenske, J. M., Likhtik, E., Russo, M. J., Gordon, J. A., Salzman, D. & Axel, R.,(2015). Neural representations of unconditioned stimuli in basolateral amygdala mediate innate and learned responses. Cell 162 (1), 134–145 https://doi.org/10.1016/j.cell.2015.06.027

Gothard, K. M. (2020). Multidimensional processing in the amygdala. Nature Reviews Neuroscience,21(10), 565–575. https://doi.org/10.1038/s41583-020-0350-y

Gothard, K. M., & Fuglevand, A. J. (2022). The role of the amygdala in processing social and affective touch. In Current Opinion in Behavioral Sciences (Vol. 43, pp. 46–53). Elsevier Ltd. https://doi.org/10.1016/j.cobeha.2021.08.004

Grabenhorst, F., Hernádi, I., & Schultz, W. (2012). Prediction of economic choice by primate amygdala neurons. Proceedings of the National Academy of Sciences of the United States of America,109(46), 18950–18955. https://doi.org/10.1073/pnas.1212706109

Grandi, L. C., & Ishida, H. (2015). The physiological effect of human grooming on the heart rate and the heart rate variability of laboratory non-human primates: A pilot study in male rhesus monkeys. Frontiers in Veterinary Science, 2(OCT). https://doi.org/10.3389/fvets.2015.00050

Granger, C.W.J. (1998). Extracting information from mega-panels and high-frequency data. Stat Neerl 52, 258–272. 10.1111/1467-9574.00084.

Hamid, A. A., Pettibone, J. R., Mabrouk, O. S., Hetrick, V. L., Schmidt, R., Vander Weele, C. M., Kennedy, R. T., Aragona, B. J., & Berke, J. D. (2015). Mesolimbic dopamine signals the value of work. Nature Neuroscience,19(1), 117–126. https://doi.org/10.1038/nn.4173

Handlin, L., Novembre, G., Lindholm, H., Kämpe, R., & Morrison, I. (2021). Human endogenous oxytocin and its neural correlates show adaptive responses to social touch based on recent social context. BioRxiv, Cmiv, 2021.04.08.438987. http://biorxiv.org/cgi/content/short/2021.04.08.438987v1?rss=1&utm_source=researcher_app&utm_medium=referral&utm_campaign=RESR_MRKT_Researcher_inbound%0Ahttps://www.biorxiv.org/content/10.1101/2021.04.08.438987v1?rss=1&utm_source=researcher_app&utm_medium

Hariri, R. & Whalen, P. (2011) The amygdala: Inside and out F1000 biology reports 3(2) http://f1000.com/reports/b/3/2

He, Q., Versteeg, C. S., Suresh, A. K., Rosenow, J., Miller, L. E., & Bensmaia, S. J. (2021). Modulation of cutaneous responses in the cuneate nucleus of macaques during active movement. bioRxiv. https://doi.org/10.1101/2021.11.15.468735

Hertenstein, M. J., Verkamp, J. M., Kerestes, A. M., & Holmes, R. M. (2007). The Communicative Functions of Touch in Humans, Nonhuman Primates, and Rats: A Review and Synthesis of the Empirical Research. 10.3200/mono.132.1.5-94

Histed, M. H., Pasupathy, A., & Miller, E. K. (2009). Learning substrates in the primate prefrontal cortex and striatum: sustained activity related to successful actions. Neuron,63(2), 244–253.

Hu B, Boyle CA & Lei S (2020). Oxytocin receptors excite lateral nucleus of central amygdala by phospholipase Cβ- and protein kinase C-dependent depression of inwardly rectifying K+ channels. J Physiology 598: 3501–3520.

Hung LW, Neuner S, Polepalli JS, Beier KT, Wright M, Walsh JJ, Lewis EM, Luo L, Deisseroth K, Dölen G & Malenka RC (2017). Gating of social reward by oxytocin in the ventral tegmental area. Science 357: 1406–1411.

Jablonski, N. G. (2021). Social and affective touch in primates and its role in the evolution of social cohesion. In Neuroscience (Vol. 464, pp. 117–125). Elsevier Ltd. https://doi.org/10.1016/j.neuroscience.2020.11.024

John, Y. J., Zikopoulos, B., Bullock, D., & Barbas, H. (2016). The Emotional Gatekeeper: A Computational Model of Attentional Selection and Suppression through the Pathway from the Amygdala to the Inhibitory Thalamic Reticular Nucleus. PLoS Computational Biology,12(2). https://doi.org/10.1371/journal.pcbi.1004722

Jovanovic, V., Fishbein, A. R., de la Mothe, L., Lee, K. F., & Miller, C. T. (2022). Behavioral context affects social signal representations within single primate prefrontal cortex neurons. Neuron. https://doi.org/10.1016/j.neuron.2022.01.020

Kaas, J. H. (1983). What, if anything, is SI? Organization of first somatosensory area of cortex. Physiological Reviews,63(1), 206–231. 10.1152/physrev.1983.63.1.206

Krzywinski, M., and Altman, N. (2014). Points of significance: Comparing samples - part II. Nature Methods 11, 355–356.

Kyriazi, P., Headley, D. B., & Pare, D. (2018). Multi-dimensional Coding by Basolateral Amygdala Neurons. Neuron,99(6), 1315-1328.e5. https://doi.org/10.1016/j.neuron.2018.07.036

Lakens, D. Calculating and reporting effect sizes to facilitate cumulative science: a practical primer for t-tests and ANOVAs. Front Psychol 4, 863 (2013).

LeDoux, J. E. (1992). Emotion and the amygdala. In J. P. Aggleton (Ed.), The amygdala: Neurobiological aspects of emotion, memory, and mental dysfunction (pp. 339–351). Wiley-Liss.

LeDoux, J. (2007). The amygdala. Current Biology,17(20). 10.1016/j.cub.2007.08.005

Lee, S. C., Amir, A., Haufler, D., & Pare, D. (2017). Differential Recruitment of Competing Valence-Related Amygdala Networks during Anxiety. Neuron,96(1), 81-88.e5. https://doi.org/10.1016/j.neuron.2017.09.002

Lehmann, J., Korstjens, A. H., & Dunbar, R. I. M. (2007). Group size, grooming and social cohesion in primates. Animal Behaviour,74(6), 1617–1629. https://doi.org/10.1016/j.anbehav.2006.10.025

Lin, M., Lucas, H.C., and Shmueli, G. (2013). Research Commentary —Too Big to Fail: Large Samples and the p-Value Problem. Inform Syst Res 24, 906–917. 10.1287/isre.2013.0480.

Livneh, U., Resnik, J., Shohat, Y., & Paz, R. (2012). Self-monitoring of social facial expressions in the primate amygdala and cingulate cortex. Proceedings of the National Academy of Sciences of the United States of America,109(46), 18956–18961. https://doi.org/10.1073/pnas.1207662109

Livneh, Y., Sugden, A. U., Madara, J. C., Essner, R. A., Flores, V. I., Sugden, L. A., Resch, J. M., Lowell, B. B., & Andermann, M. L. (2020) Estimation of current and future physiological state in Insular cortex. Neuron, 105(6), 1094-1111.e10. https://doi.org/10.1016/j.neuron.2019.12.027

Löken, L. S., Evert, M., & Wessberg, J. (2011). Pleasantness of touch in human glabrous and hairy skin: Order effects on affective ratings. Brain Research, 1417, 9–15. https://doi.org/10.1016/j.brainres.2011.08.011

Lucas, M. v., Anderson, L. C., Bolling, D. Z., Pelphrey, K. A., & Kaiser, M. D. (2015). Dissociating the neural correlates of experiencing and imagining affective touch. Cerebral Cortex,25(9), 2623–2630. https://doi.org/10.1093/cercor/bhu061

Mackes, N. K., Golm, D., Sarkar, S., Kumsta, R., Rutter, M., Fairchild, G., Mehta, M. A., & Sonuga-Barke, E. J. S. (2020). Early childhood deprivation is associated with alterations in adult brain structure despite subsequent environmental enrichment. PNAS. https://doi.org/10.1073/pnas.1911264116/-/DCSupplemental

Mante, V., Sussillo, D., Shenoy, K. v., & Newsome, W. T. (2013). Context-dependent computation by recurrent dynamics in prefrontal cortex. Nature,503(7474), 78–84. https://doi.org/10.1038/nature12742

McFarland, R., & Majolo, B. (2011). Grooming coercion and the post-conflict trading of social services in wild Barbary macaques. PLoS ONE,6(10). https://doi.org/10.1371/journal.pone.0026893

McGlone, F., Wessberg, J., & Olausson, H. (2014). Discriminative and Affective Touch: Sensing and Feeling. In Neuron (Vol. 82, Issue 4, pp. 737–755). Cell Press. https://doi.org/10.1016/j.neuron.2014.05.001

Moberg, K., & Petersson, M. (2022). Physiological effects induced by stimulation of cutaneous sensory nerves, with a focus on oxytocin. Current Opinion in Behavioral Sciences, 43, 159–166 https://doi.org/10.1016/j.cobeha.2021.10.001

Morrow, J., Mosher, C., & Gothard, K. (2019). Multisensory neurons in the primate amygdala. Journal of Neuroscience,39(19), 3663–3675. https://doi.org/10.1523/JNEUROSCI.2903-18.2019

Mosher, C. P., Zimmerman, P. E., Fuglevand, A. J., & Gothard, K. M. (2016). Tactile stimulation of the face and the production of facial expressions activate neurons in the primate amygdala. ENeuro,3(5). https://doi.org/10.1523/ENEURO.0182-16.2016

Nelson RJ, Sur M, Felleman DJ & Kaas JH (1980). Representations of the body surface in postcentral parietal cortex of Macaca fascicularis. J Comp Neurol 192: 611–643.

Nummenmaa, L., Tuominen, L., Dunbar, R., Hirvonen, J., Manninen, S., Arponen, E., Machin, A., Hari, R., Jääskeläinen, I. P., & Sams, M. (2016). Social touch modulates endogenous μ-opioid system activity in humans. NeuroImage, 138, 242–247. https://doi.org/10.1016/j.neuroimage.2016.05.063

Nuzzo, R. (2014). Scientific method: Statistical errors. Nature 506, 150–152. 10.1038/506150a.

O’Connor TG, Rutter M: Attachment disorder behavior following early severe deprivation: extension and longitudinal follow-up. English and Romanian Adoptees Study Team. J Am Acad Child Adolesc Psychiatry (2000), 39:703–712. 10.1097/00004583-200006000-00008

Olausson, H., Wessberg, J., Morrison, I., McGlone, F., and Vallbo, A. (2010). The neurophysiology of unmyelinated tactile afferents. Neuroscience & Biobehavioral Reviews 34, 185–191. 10.1016/j.neubiorev.2008.09.011.

Porges, S. W. (2001). The polyvagal theory: phylogenetic substrates of a social nervous system. International Journal of Psychophysiology, 42, 123–146. https://doi.org/10.1016/S0167-8760(01)00162-3

Putnam, P. & Gothard, K. M., (2019). Multi-dimensional neural selectivity in the primate amygdala. eNeuro 10.1523/ENEURO.0153-19.2019

Rainer, G., Asaad, W. F., & Miller, E. K. (1998). Selective representation of relevant information by neurons in the primate prefrontal cortex. Nature. https://doi.org/10.1038/31235

Resnik, J. & Paz, R. (2015). Fear generalization in the primate amygdala. Nature Neuroscience, 18(2), 188–190 https://doi.org/10.1038/nn.3900

Rolls, E. T., O’Doherty, J., Kringelbatch, M. L., Francis, S., Bowtell, R., & McGlone, F. (2003). Representations of Pleasant and Painful Touch in the Human Orbitofrontal and Cingulate Cortices. Cerebral Cortex, 13(3), 308–317. https://doi.org/10.1093/cercor/13.3.308

Rutter, M. (1998). Developmental Catch-up, and Deficit, Following Adoption after Severe Global Early Privation. In J. Child Psychol. Psychiat (Vol. 39, Issue 4). https://doi.org/10.1111/1469-7610.00343

Saez, A., Rigotti, M., Ostojic, S., Fusi, S., & Salzman, C. D. (2015). Abstract Context Representations in Primate Amygdala and Prefrontal Cortex. Neuron,87(4), 869–881. https://doi.org/10.1016/j.neuron.2015.07.024

Sanchez, M. M. (2006). The impact of early adverse care on HPA axis development: Nonhuman primate models. Hormones and Behavior,50(4), 623–631. https://doi.org/10.1016/j.yhbeh.2006.06.012

Schino, G., & Aureli, F. (2008). Grooming reciprocation among female primates: A meta-analysis. Biology Letters,4(1), 9–11. https://doi.org/10.1098/rsbl.2007.0506

Schino, G., Scucchi, S., Maestripieri, D., & Turillazzi, P. G. (1988). Allogrooming as a Tension-Reduction Mechanism: A Behavioral Approach. In American Journal of Primatology (Vol. 16). 10.1002/ajp.1350160106

Schultz, W., Dayan, P., & Montague, P. R. (1997). A neural substrate of prediction and reward. Science,275(5306), 1593–1599. https://doi.org/10.1126/science.275.5306.1593

Spitler, K. M., & Gothard, K. M. (2008). A removable silicone elastomer seal reduces granulation tissue growth and maintains the sterility of recording chambers for primate neurophysiology. Journal of Neuroscience Methods,169(1), 23–26.https://doi.org/10.1016/j.jneumeth.2007.11.026

Steinmetz, N. A., Zatka-Haas, P., Carandini, M., & Harris, K. D. (2019). Distributed coding of choice, action and engagement across the mouse brain. Nature,576(7786), 266–273. https://doi.org/10.1038/s41586-019-1787-x

Suvilehto, J. T., Renvall, V., & Nummenmaa, L. (2021). Relationship-specific Encoding of Social Touch in Somatosensory and Insular Cortices. Neuroscience, 464, 105–116. https://doi.org/10.1016/j.neuroscience.2020.09.015

Taira’ And, K., & Rolls, E. T. (1996). Receiving Grooming as a Reinforcer for the Monkey. In Physiology & Behavior (Vol. 59, Issue 6). https://doi.org/10.1016/0031-9384(95)02213-9

Tang, Y., Benusiglio, D., Lefevre, A., Hilfiger, L., Althammer, F., Bludau, A., … & Grinevich, V. (2020). Social touch promotes interfemale communication via activation of parvocellular oxytocin neurons. Nature neuroscience,23(9), 1125–1137. https://doi.org/10.1038/s41593-020-0674-y

Taub, A. H., Shohat, Y., & Paz, R. (2018). Long time-scales in primate amygdala neurons support aversive learning. Nature Communications,9(1). https://doi.org/10.1038/s41467-018-07020-4

Tiddi, B., Aureli, F., & Schino, G. (2012). Grooming up the hierarchy: The exchange of grooming and rank-related benefits in a new world primate. PLoS ONE,7(5). https://doi.org/10.1371/journal.pone.0036641

Triscoli, C., Croy, I., Olausson, H., & Sailer, U. (2017). Touch between romantic partners: Being stroked is more pleasant than stroking and decelerates heart rate. Physiology and Behavior, 177, 169–175. https://doi.org/10.1016/j.physbeh.2017.05.006

Turchi, J., Chang, C., Ye, F. Q., Russ, Be. E., Yu, D. K., Cortex, C. R., Monosov, I. E., Duyn, J. H., & Leopold, D. A. (2018). The basal forebrain regulates global resting-state fMRI fluctuations. Neuron, 97(4), 940952.e4. https://doi.org/10.1016

Tsodyks, M., Kenet, T., Grinvald, A., & Arieli, A. (1999). Linking Spontaneous Activity of Single Cortical Neurons and the Underlying Functional Architecture. 10.1126/science.286.5446.1943

Tye, K. M., & Janak, P. H. (2007). Amygdala neurons differentially encode motivation and reinforcement. Journal of Neuroscience,27(15), 3937–3945. https://doi.org/10.1523/JNEUROSCI.5281-06.2007

Uvnäs Moberg, K., & Petersson, M. (2022). Physiological effects induced by stimulation of cutaneous sensory nerves, with a focus on oxytocin. In Current Opinion in Behavioral Sciences (Vol. 43, pp. 159–166). Elsevier Ltd. https://doi.org/10.1016/j.cobeha.2021.10.001

Vallbo, A.B., Olausson, H., Wessberg, J., and Kakuda, N. (1995). Receptive field characteristics of tactile units with myelinated afferents in hairy skin of human subjects. J Physiology 483, 783–795. 10.1113/jphysiol.1995.sp020622.

von Mohr, M., Kirsch, L. P., & Fotopoulou, A. (2017). The soothing function of touch: Affective touch reduces feelings of social exclusion. Scientific Reports,7(1). https://doi.org/10.1038/s41598-017-13355-7

Walker, S. C., Cavieres, A., Peñaloza-Sancho, V., El-Deredy, W., McGlone, F. P., & Dagnino-Subiabre, A. (2020). C-low threshold mechanoafferent targeted dynamic touch modulates stress resilience in rats exposed to chronic mild stress. European Journal of Neuroscience. https://doi.org/10.1111/ejn.14951

Wang, S., Yu, R., Tyszka, J. M., Zhen, S., Kovach, C., Sun, S., Huang, Y., Hurlemann, R., Ross, I. B., Chung, J. M., Mamelak, A. N., Adolphs, R., & Rutishauser, U. (2017). The human amygdala parametrically encodes the intensity of specific facial emotions and their categorical ambiguity. Nature Communications, 8. https://doi.org/10.1038/ncomms14821

